# Keratin 17 Defines an Epithelial-Mesenchymal Transition-Associated Migratory Cell State in Pancreatic Ductal Adenocarcinoma

**DOI:** 10.64898/2026.10.27.762113

**Authors:** Shayan Sarkar, Monisankar Ghosh, Lyanne Delgado-Coka, Scott Powers, Rekha Sathian, Ramana Davuluri, Xiangyu Gong, Michael Mak, Jorge Villar, Xiaohua Qi, Sruthi Babu, Natalia Marchenko, Kenneth R. Shroyer

**Author notes:** Corresponding author: Kenneth R. Shroyer, SUNY Distinguished Professor, The Marvin Kuschner Professor and Chair of Pathology, Basic Science Tower level 9, Rm 142, Renaissance School of Medicine, Stony Brook University, Stony Brook, NY 11794-8691. Senior co-authors (Natalia Marchenko, Kenneth R. Shroyer).

## Abstract

Pancreatic ductal adenocarcinoma (PDAC) is characterized by early invasion, metastasis, and therapeutic resistance that contribute substantially to its poor clinical outcome. Keratin 17 (K17) is a recognized marker of the aggressive basal-like subtype of PDAC and an independent predictor of adverse prognosis; however, the biological basis for this association remains incompletely understood. To define K17-associated cellular programs, we integrated single-cell RNA sequencing (scRNA-seq) data from 42 patients, Clinical Proteomic Tumor Analysis Consortium (CPTAC) proteomic data from 140 tumors, MACSima multiplex spatial imaging of 63 patients, dual-plex immunohistochemistry of 60 cases, and three-dimensional spheroid migration assays. Single-cell transcriptomic analysis demonstrated significant enrichment of epithelial-mesenchymal transition (EMT)-associated gene programs among K17-positive malignant epithelial cells. Consistent with this finding, K17-positive tumors in the CPTAC cohort exhibited enrichment of the Hallmark EMT pathway at the protein level. Among six candidate markers identified through transcriptomic analyses, laminin subunit gamma-2 (LAMC2) was the only candidate associated with K17 at every validation step. Tumors positive for both K17 and LAMC2 had significantly worse overall survival than double-negative tumors (adjusted hazard ratio = 2.53). Multiplex spatial imaging localized K17-positive cells predominantly to poorly differentiated tumor regions and demonstrated frequent single-cell co-expression with LAMC2. Whole-slide dual-plex immunohistochemistry further confirmed the association between K17 and LAMC2 and showed progressive enrichment of K17/LAMC2-positive tumor cells with increasing intratumoral histologic grade. Functionally, gain- and loss-of-function studies in three-dimensional PDAC spheroids demonstrated that K17 promotes migratory outgrowth, and differences in proliferation did not account for this effect. Collectively, these findings identify K17 as a marker of a poorly differentiated, EMT-associated migratory cell state in PDAC and provide functional evidence that K17 contributes to tumor-cell migration.

## Introduction

Pancreatic ductal adenocarcinoma (PDAC) remains one of the deadliest human malignancies, with a 5-year survival rate of approximately 13%.^1^ Despite advances in surgical and systemic therapies, patient outcomes remain limited due to the propensity of PDAC for early local invasion, metastatic dissemination, and therapeutic resistance. Experimental evidence suggests that pancreatic epithelial cells can disseminate before the development of overt malignant lesions in mouse models, underscoring the intrinsically aggressive nature of this disease.^2^ In addition, epithelial-mesenchymal transition (EMT) and tumor cell plasticity have been implicated in metastatic progression and resistance to conventional therapies in PDAC.^2,3^ A more comprehensive understanding of the biological mechanisms underlying PDAC progression is therefore needed to identify clinically actionable vulnerabilities.

Keratin 17 (K17) has been regarded as one of the most robust biomarkers of adverse prognosis in PDAC. Its expression is associated with the basal-like molecular subtype and independently predicts poor patient survival.^4^ The use of K17 as an immunohistochemical biomarker was first developed by our group in cervical squamous lesions.^5^ Beyond its prognostic value, K17 has been implicated in multiple cancer-promoting processes, including enhanced tumor growth, therapeutic resistance, immune modulation, and cellular plasticity.^6^ More recently, we reported that K17 expression is associated with distinct immune cell topography in PDAC^7^ and promotes chemoresistance through metabolic reprogramming.^8^ Collectively, these findings suggest that K17 functions as more than a passive biomarker and may actively contribute to biological pathways that drive disease progression.

Molecular classification studies have identified basal-like (also termed quasi-mesenchymal or squamous) tumors as having worse survival than classical tumors,^9–11^ and basal-like states have been linked to cellular plasticity and differential drug response.^12^ More recent transcriptomic, spatial, and single-cell studies have refined this paradigm by demonstrating that PDAC subtypes exist along a continuum of dynamic cellular states rather than as discrete biological entities. These studies have revealed substantial intratumoral heterogeneity, including the coexistence of basal-like and classical programs within individual tumors.^13,14^ Nevertheless, enrichment of basal-like programs remains consistently linked to aggressive disease biology and inferior clinical outcomes. Given its association with this cellular state, K17 represents a particularly compelling candidate for investigating the mechanisms that underlie aggressive PDAC behavior.

Despite this well-established association with poor clinical outcome, the programs through which K17 is linked to aggressive disease remain unclear. Defining the molecular and cellular programs associated with K17 expression may provide important insight into how K17 relates to PDAC progression. To address this, we performed a multimodal investigation beginning with single-cell transcriptomic profiling of malignant epithelial cells. Single-cell approaches provide a powerful framework for identifying biologically distinct tumor populations that may be obscured by bulk analyses and are therefore well suited to define K17-associated cellular programs. We subsequently integrated proteomic, spatial, histologic, and functional analyses to evaluate the reproducibility of K17-associated phenotypes across platforms and to determine whether K17 influences tumor-cell behavior.

Using this integrated approach, we identify K17 as a marker of an EMT-associated migratory tumor-cell state that preferentially localizes to poorly differentiated regions of PDAC. We further identify laminin subunit gamma-2 (LAMC2), a protein implicated in invasion and EMT-related biology, as a reproducible correlate of this phenotype across transcriptomic, proteomic, spatial, and histologic analyses. Finally, we show that K17 expression increases migratory outgrowth in three-dimensional PDAC spheroid models. Together, these findings characterize an EMT-associated migratory state linked to K17 and establish a foundation for mechanistic studies of K17 in PDAC invasion and dissemination.

## Materials and Methods

### Computational Analysis of Single-Cell RNA Sequencing (scRNA-seq) Data

An integrated scRNA-seq dataset of PDAC tumors was assembled from two publicly available sources.^15,16^ Raw sequencing data were processed into count matrices using CellRanger (version 7.1.0).^17^ Ambient RNA contamination was removed using CellBender remove-background, a deep generative Bayesian model that infers and subtracts ambient RNA while preserving biological signal.^18^

Following quality-control filtering, the final integrated dataset comprised cells from 42 PDAC patients (22 from Peng et al. and 20 from Werba et al.). Normalization, dimensionality reduction, clustering, and cell-type annotation were performed using Scanpy.^19^ Epithelial cell populations were identified based on *KRT19* expression (Figure 1B). Malignant epithelial cells were subsequently distinguished from benign epithelial cells using inferCNVpy-based copy-number variation inference, with immune, endothelial, and stromal populations serving as diploid reference cells. Cells exhibiting recurrent chromosome-scale copy-number alterations were classified as malignant, yielding 77,155 malignant epithelial cells. Batch effects across samples were corrected using Harmony, and missing expression values were imputed using Adaptively-thresholded Low Rank Approximation (ALRA).^20,21^

**Figure 1.**
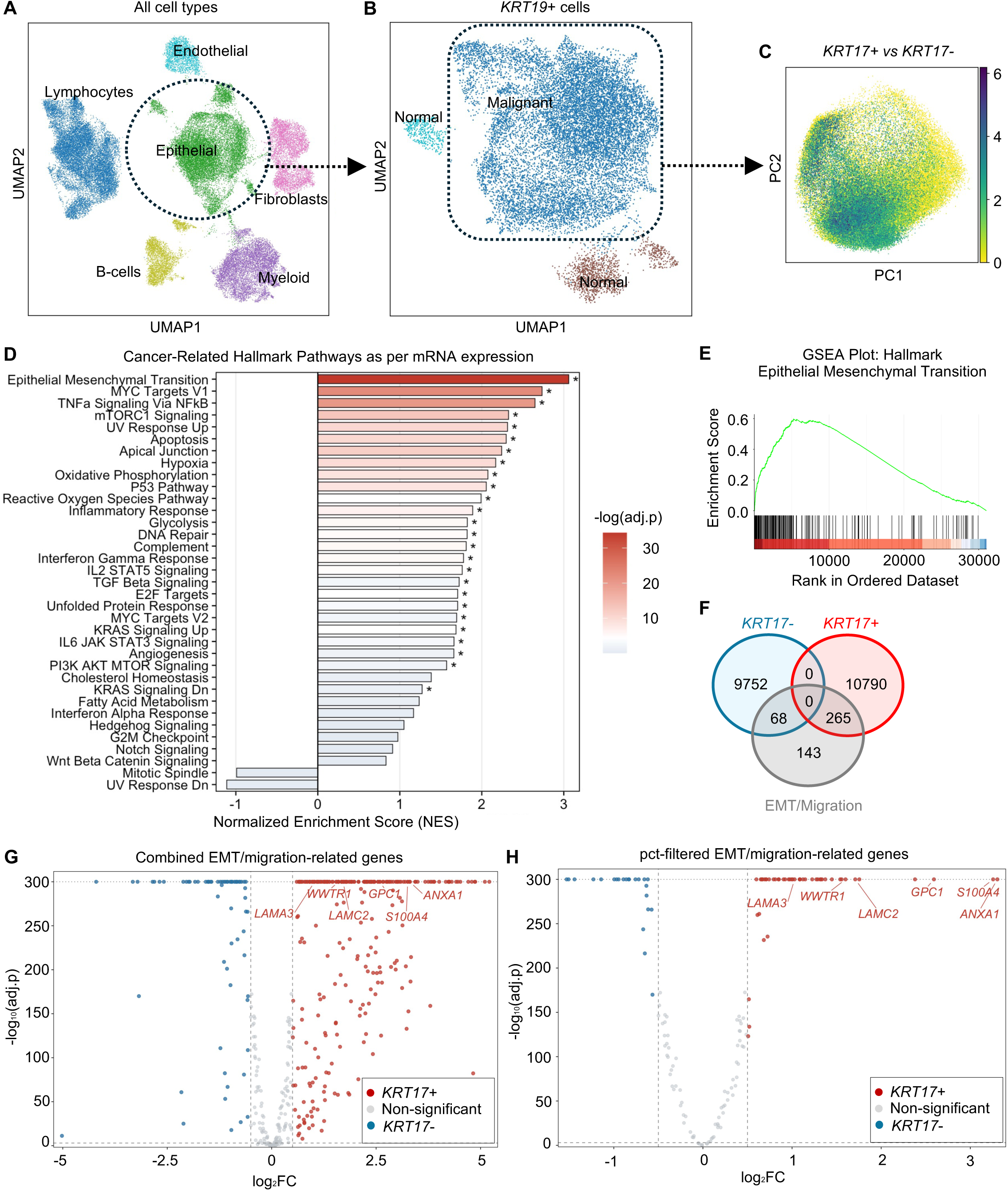
*KRT17*-positive malignant epithelial cells are enriched for EMT- and migration-associated transcriptional programs in PDAC. (A) UMAP of all captured cells, colored by major cell type. (B) Re-clustering of the epithelial compartment resolves malignant and normal populations. (C) PCA projection of malignant cells colored by continuous *KRT17* expression level, illustrating a gradient rather than a discrete bimodal distribution. (D) Hallmark pathway enrichment (normalized enrichment score, NES) in *KRT17*-positive vs. *KRT17*-negative malignant cells; EMT is the top-ranked pathway. Asterisks denote pathways with significant enrichment (adjusted p < 0.05). (E) GSEA enrichment plot for the Hallmark EMT gene set (NES = 3.06, FDR < 0.05). (F) Venn diagram of genes enriched in *KRT17*-positive cells, *KRT17*-negative cells, and the curated EMT/migration gene set, showing overlap between each group and the EMT/migration signature. (G) Volcano plot of differential EMT/migration gene expression in *KRT17*-positive vs. *KRT17*-negative malignant cells, with six selected EMT/migration markers (ANXA1, S100A4, GPC1, LAMC2, WWTR1, LAMA3) labeled. (H) Same comparison as (G) after filtering to genes with a minimum expression prevalence of 20% among malignant cells.

Expression of the *KRT17* gene was evaluated within the malignant epithelial compartment using the normalized ALRA-imputed expression matrix. Cells with detectable *KRT17* expression (*KRT17* > 0) were classified as *KRT17*-positive, whereas cells with an expression value of 0 were classified as *KRT17*-negative. For visualization, the KRT17 color scale was capped to display expression differences; this cap was not used to define KRT17-positive cells or in any analysis. Differential gene-expression analysis between *KRT17*-positive and *KRT17*-negative malignant epithelial cells was subsequently performed using Scanpy.

### Gene Set Enrichment Analysis

Gene set enrichment analysis (GSEA) was performed using fgsea in R (version 4.4.2), testing a curated panel of cancer-related Hallmark gene sets.^22^ Pathways evaluated included Epithelial-Mesenchymal Transition, TGF-β Signaling, MYC Targets, and KRAS Signaling, among other cancer-related hallmark pathways. Minimum and maximum gene set sizes were set to 15 and 500 genes, respectively. Normalized enrichment score (NES), false discovery rate (FDR)-adjusted p-value, and leading-edge gene ratio were reported for each pathway.

To identify *KRT17*-associated EMT/migration markers, differentially expressed genes between *KRT17*-positive and *KRT17*-negative malignant cells were intersected with a curated EMT/migration gene set comprising the Hallmark EMT gene set together with the Gene Ontology Biological Process terms Epithelial to Mesenchymal Transition and Positive Regulation of Epithelial Cell Migration. Genes were subsequently filtered to a minimum expression prevalence of 20% among malignant cells. Six markers, namely Annexin A1 (ANXA1), S100 calcium binding protein A4 (S100A4), glypican-1 (GPC1), LAMC2, WW domain-containing transcription regulator 1 (WWTR1), and laminin subunit alpha-3 (LAMA3), were selected for downstream protein-level validation based on differential expression magnitude, statistical significance, prior biological relevance to EMT or PDAC, and availability of validated antibodies for tissue-based assays.

### CPTAC Proteomic Analysis

Proteomic data from the Clinical Proteomic Tumor Analysis Consortium (CPTAC) PDAC cohort (n = 140 treatment-naive tumors) were obtained from the LinkedOmics data portal, including the tumor proteome matrix and clinical annotation file.^23^ Analyses were performed in R (version 4.4.2); cohort characteristics are summarized in Table 1. K17 and LAMC2 protein abundance were correlated using Spearman rank correlation, selected given the non-parametric distribution characteristic of tandem mass tag (TMT) proteomics data. For differential protein abundance analysis, samples were stratified by median K17 abundance into K17-positive and K17-negative groups (n = 70 each); proteins quantified in at least 50% of samples per group were retained (8,665 proteins); missing values were imputed by row minimum, and differential abundance was assessed using limma (version 3.62.2) with empirical Bayes moderation and Benjamini-Hochberg correction (FDR < 0.05, |log2 fold change| > 0.5).^24^ GSEA was performed on the resulting ranked t-statistics against the Hallmark gene sets (including the Hallmark EMT set; 158 of 200 genes quantified).

**Table 1.** Demographic and Clinicopathologic Characteristics of the CPTAC PDAC Cohort.

| **Characteristic** | **Full Cohort (n = 140)** | **Survival Analysis Subset (n = 123)** |
| --- | --- | --- |
| Age, years, median (range) | 65 (31–85) | 64 (31–85) |
| **Sex** |  |  |
| Male | 74 (52.9%) | 65 (52.8%) |
| Female | 66 (47.1%) | 58 (47.2%) |
| **Race** |  |  |
| Not reported | 107 (76.4%) | 96 (78.0%) |
| White | 31 (22.1%) | 25 (20.3%) |
| Black or African American | 2 (1.4%) | 2 (1.6%) |
| **Tumor Site** |  |  |
| Head | 103 (74.6%) | 94 (76.4%) |
| Tail | 14 (10.1%) | 11 (8.9%) |
| Body | 14 (10.1%) | 13 (10.6%) |
| Body and tail | 5 (3.6%) | 4 (3.3%) |
| Head and body | 2 (1.4%) | 1 (0.8%) |
| Missing | 2 | 0 |
| **AJCC Pathological Stage** |  |  |
| Stage IIB | 52 (38.8%) | 48 (39.0%) |
| Stage III | 42 (31.3%) | 37 (30.1%) |
| Stage IB | 16 (11.9%) | 15 (12.2%) |
| Stage IV | 9 (6.7%) | 9 (7.3%) |
| Stage IIA | 8 (6.0%) | 7 (5.7%) |
| Stage IA | 7 (5.2%) | 7 (5.7%) |
| Missing | 6 | 0 |
| **Primary Tumor (pT)** |  |  |
| pT2 | 86 (61.4%) | 75 (61.0%) |
| pT3 | 41 (29.3%) | 37 (30.1%) |
| pT1c | 9 (6.4%) | 9 (7.3%) |
| pTX | 2 (1.4%) | 0 (0%) |
| pT4 | 1 (0.7%) | 1 (0.8%) |
| pT1a | 1 (0.7%) | 1 (0.8%) |
| **Regional Lymph Nodes (pN)** |  |  |
| pN1 | 54 (38.6%) | 50 (40.7%) |
| pN2 | 47 (33.6%) | 42 (34.1%) |
| pN0 | 31 (22.1%) | 29 (23.6%) |
| pNX | 8 (5.7%) | 2 (1.6%) |
| **Residual Tumor Status** |  |  |
| R0: No residual tumor | 73 (52.9%) | 66 (53.7%) |
| R1: Microscopic residual tumor | 34 (24.6%) | 33 (26.8%) |
| RX: Cannot be assessed | 25 (18.1%) | 19 (15.4%) |
| R2: Macroscopic residual tumor | 6 (4.3%) | 5 (4.1%) |
| Missing | 2 | 0 |
| **Tumor Focality** |  |  |
| Unifocal | 135 (97.8%) | 121 (98.4%) |
| Multifocal | 3 (2.2%) | 2 (1.6%) |
| Missing | 2 | 0 |
| **Vital Status** |  |  |
| Deceased | 77 (56.6%) | 69 (56.1%) |
| Living | 59 (43.4%) | 54 (43.9%) |
| Missing | 4 | 0 |

### Survival Analysis

Clinical outcome data were available for 123 of 140 patients. K17 and LAMC2 were dichotomized separately according to analytic goal: a median split was used for differential abundance analysis and GSEA, whereas survival analysis used protein-specific optimal cut-points derived in SAS (version 9.4), selected to identify the abundance threshold most strongly associated with outcome. Patients were classified by combined K17/LAMC2 status into four groups, and survival was estimated using Kaplan-Meier curves and compared by log-rank test. Cox proportional hazards models estimated hazard ratios (HR) for K17 and LAMC2 individually and jointly, adjusted for tumor stage and age at diagnosis.

### Case Selection for MACSima and IHC

Formalin-fixed, paraffin-embedded archival tissue blocks of human PDACs were provided by the Biobank (IRB ID: 94651_MODCR016) of the Renaissance School of Medicine (RSOM) Cancer Center Biobank Core Laboratory to create two overlapping cohorts for MACSima and dual-plex immunohistochemistry (IHC) studies. The MACSima cohort comprised 69 tissue cores, obtained from 63 unique Stony Brook University Hospital patients with primary PDAC, selected based on hematoxylin and eosin (H&E) evaluation (cohort characteristics in Supplementary Table S1); six patients each contributed two cores to the tissue microarray (TMA). The dual-plex IHC cohort comprised 60 FFPE specimens selected by H&E examination of representative sections, independent of K17 or LAMC2 status (Table 2). Tumor stage (American Joint Committee on Cancer or AJCC 8th edition) and whole-tumor histopathologic grade (World Health Organization or WHO) were obtained from the surgical pathology reports.^25–27^

**Table 2.** Demographic and Clinicopathologic Characteristics of the Dual-Plex IHC Cohort.

| **Characteristic** | **n** | **%** |
| --- | --- | --- |
| Total patients | 60 |  |
| **Age at diagnosis, years** |  |  |
| Mean ± SD | 68.9 ± 10.4 | |
| Median (range) | 70.5 (43–89) |  |
| **Sex** |  |  |
| Male | 35 | 58.3% |
| Female | 25 | 41.7% |
| **Race** |  |  |
| White | 55 | 91.7% |
| Other | 4 | 6.7% |
| Black | 1 | 1.7% |
| **Vital status** |  |  |
| Dead | 40 | 66.7% |
| Alive | 20 | 33.3% |
| **Pathologic AJCC stage group** |  |  |
| Stage IB | 4 | 6.7% |
| Stage IIA | 4 | 6.7% |
| Stage IIB | 38 | 63.3% |
| Stage III | 6 | 10.0% |
| Stage IV | 2 | 3.3% |
| Not available | 6 | 10.0% |
| **Chemotherapy** |  |  |
| Yes | 46 | 76.7% |
| No | 10 | 16.7% |
| Not available | 4 | 6.7% |

### Histologic Grading of Individual Morphologic Components

Intratumoral morphologic heterogeneity was characterized by classifying discrete regions within each tumor as grade I (well-differentiated, gland-forming, preserved apicobasal polarity), grade II (moderately differentiated, partial loss of glandular architecture and nuclear polarity), or grade III (solid sheets, trabeculae, or small clusters and single cells that lacked evidence of glandular differentiation). Grading was performed by a board-certified surgical pathologist and is distinct from the overall tumor grade assigned during case selection; a single case may contain regions of more than one intratumoral grade (Figure 5G). This grading scheme is described in detail in a related manuscript from our group (Delgado-Coka et al., manuscript under review, *Scientific Reports*).

### Multiplex Imaging (MACSima)

Each TMA block contained approximately 13 cores, each 4 mm in diameter. Multiplex spatial protein analysis was performed on 4 µm thick FFPE tissue sections using the MACSima™ Imaging Platform (Miltenyi Biotec), with automated cyclic staining, imaging, and signal erasure (Supplementary Figure S3A). The antibody panel is presented in Supplementary Table S2. It included six K17-associated EMT/migration markers identified through transcriptomic analysis together with K17 and pan-cytokeratin (PanCK; low-molecular-weight cytokeratins 7, 8, 18 and 19) for tumor-cell identification.

### Image Segmentation and Cell Population Gating and Quantification

Images were processed using MACS-iQ View software (Miltenyi Biotec) to evaluate the spatial relationship between K17 and the EMT-associated marker panel. Tumor regions of interest (ROIs) were annotated based on tissue morphology, excluding pancreatitis and pancreatic intraepithelial neoplasia to restrict analyses to PDAC cells. The parameters for segmentation and the results are shown in Supplementary Table S3 and Supplementary Figure S3B-C.

Tumor cells were identified using an inclusive (OR) gating strategy based on expression of PanCK and K17; marker-specific intensity thresholds were determined, supported by visual examination. K17-positive tumor cells were defined by an intensity threshold gate, with K17-negative tumor cells obtained by Boolean complement within the tumor-cell gate. Marker positivity within each group was quantified using thresholds applied consistently across cores within a given TMA.

### Marker Panel Structure and Handling of Staining Failure

The final panel comprised six K17-associated EMT/migration markers, K17, PanCK, and DAPI (Supplementary Figure S3D-L). Cluster-level analyses (Figure 4A-C) were restricted to tissue cores with successful staining across all markers (N = 66 of 69 cores). Three cores were excluded because of marker-specific staining failure (two LAMC2 and one S100A4). The box-plot analyses used all available cores with successful staining for the marker being assessed (LAMC2, N = 67; ANXA1, N = 69; S100A4, N = 68; GPC1, WWTR1, and LAMA3, N = 69 each).

### Data Processing and Clustering Analysis

Single-cell marker intensity measurements from ROIs across TMA slides were pooled for analysis. Markers and ROI-specific measurements with unreliable staining or substantial missingness were excluded. Only cells with complete measurements across all markers were retained for downstream dimensionality reduction and clustering.

Marker intensities were transformed using the inverse hyperbolic sine (arcsinh) function with a cofactor of 150. To mitigate systematic differences in intensity distributions between TMA slides, marker-specific Z-score standardization was performed independently within each TMA. The resulting values were pooled and globally standardized prior to dimensionality reduction. Principal component analysis (PCA) was performed on the standardized marker matrix, and a nearest-neighbor graph was constructed using 30 neighbors in PCA space. Cells were clustered using the Leiden algorithm.^28^ Uniform Manifold Approximation and Projection (UMAP) was applied to the standardized marker matrix for two-dimensional visualization.

### Immunohistochemistry

Single-plex IHC for K17 and LAMC2 was performed on separate FFPE sections. Following deparaffinization and antigen retrieval (Citric Acid Buffer, Vector Laboratories, ref: H-3300; 120°C for 10 minutes), endogenous peroxidase was blocked with hydrogen peroxide (Sigma-Aldrich, CAS: 7722-84-1). Sections were incubated with anti-K17 antibody (clone D12E5, Cell Signaling Technology, #12509, 1:800 dilution) or anti-LAMC2 antibody (clone E-6, Santa Cruz Biotechnology, sc-28330, 1:2000 dilution), detected using the VECTASTAIN Elite ABC-HRP Kit (Vector Laboratories, ref: PK-7200) and visualized with 3,3′-diaminobenzidine or DAB (ref: SK-4103-100) chromogen. Slides were counterstained with hematoxylin; representative images are shown in Supplementary Figure S5. The same protocol was used to stain sections of L3.6 spheroids (gain-of-function model) with the K17 antibody (Supplementary Figure S6D-E).

Dual-plex IHC for K17 and LAMC2 followed identical protocols for deparaffinization, antigen retrieval, and biotin blocking. LAMC2 was stained first and visualized with DAB; K17 was subsequently stained and visualized with Vina Green chromogen (Biocare Medical, ref: BRR807AS). Slides were counterstained with hematoxylin and mounted using a resin-based medium.

### Cell Culture

The human L3.6 pancreatic adenosquamous cell line (KRAS^G12A^), which expresses endogenous K17, was provided by Dr. Wei-Xing Zong (Rutgers University). PANC-1 human PDAC cells (CRL-1469, KRAS^G12D^, TP53^R273H^, RRID: CVCL_0480) were obtained from the American Type Culture Collection (ATCC). Cells were cultured in Dulbecco’s Modified Eagle Medium (DMEM, Gibco, #11995-065) supplemented with 10% fetal bovine serum (FBS, NEUROMICS, #FBS002) and 1% penicillin-streptomycin (Gibco, #15140-122). Cells were routinely tested for Mycoplasma contamination and were used within 10 passages of thawing.

To assess the functional contribution of K17 to tumor-cell migration, isogenic K17 loss- and gain-of-function PDAC models were generated in L3.6 cells, a human pancreatic cancer cell line widely used as a model of aggressive/metastatic PDAC. K17 knockout cells were generated using CRISPR-Cas9-mediated genome editing, whereas K17-overexpressing cells were generated through stable ectopic expression of human K17.^29,30^ Empty-vector-transduced cells served as controls for overexpression studies. Generation and validation of K17-modified cell models followed approaches previously established by our group.^8^ PANC-1 cells were used as an independent gain-of-function model to confirm findings across a second PDAC cell line.

### 3D Spheroid Assay

Three-dimensional spheroid migration assays were designed to evaluate migratory behavior in a physiologically relevant extracellular matrix context, adapting the collagen-interface invasion model in which spheroids migrate along the interface beneath a three-dimensional collagen layer.^31^ Briefly, 1,000 cells were seeded per well of a 96-well low-attachment U-bottom plate (Corning, cat: 07-200-720A) and allowed to self-aggregate into spheroids for 3 days. They were subsequently embedded in neutralized collagen I (Corning, CLS354249) at 2 mg/mL and dispensed as domes at the matrix-surface interface (3-5 spheroids per dome, approximately). Following polymerization, complete medium was added, and spheroids were maintained for the indicated duration, during which tumor cells migrated outward from the spheroid core into the surrounding collagen matrix. Phase-contrast images were acquired at defined time points, and total spheroid and core spheroid areas were measured using ImageJ (Fiji).^32^

### Migration Index

Migratory outgrowth was quantified using a normalized migration index calculated at the experimental endpoint (96 hours for L3.6 and 48 hours for PANC-1):

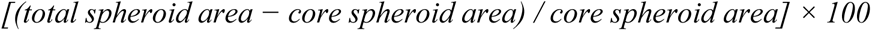

This normalization accounts for variation in initial spheroid size and allows comparison of migratory behavior across experiments.

### Confocal Imaging

Spheroids were fixed in 4% paraformaldehyde, permeabilized in Triton X-100, and blocked in bovine serum albumin. They were incubated with anti-K17 (Cell Signaling Technology, #12509) or anti-Ki-67 (Cell Signaling Technology, #9129, RRID:AB_2687446) primary antibody, followed by the corresponding fluorophore-conjugated secondary antibody and Hoechst 33342 for nuclear counterstaining. Imaging was performed using a Leica SP8 confocal microscope, with images processed in ImageJ (Fiji).

### Ki-67 Index Quantification

The Ki-67 proliferative index was calculated as the ratio of mean Ki-67 to mean Hoechst fluorescence intensity per spheroid, normalizing for variation in nuclear density.

### Statistical Analysis

Associations between K17 and candidate markers were assessed by Spearman rank correlation (single-cell, CPTAC, and cluster-level analyses). Survival was analyzed by Kaplan-Meier curves with log-rank tests and Cox proportional hazards models adjusted for stage and age; these analyses were considered exploratory. MACSima marker positivity in K17-positive versus K17-negative tumor cells was compared across cores by Wilcoxon matched-pairs signed-rank test. Differences in K17/LAMC2 category composition across intratumoral grades (Figure 5H–J) were tested by Friedman test, with post hoc Wilcoxon signed-rank tests and a single Benjamini-Hochberg correction across all 12 comparisons (rstatix, version 0.7.3).

Spheroid experiments were performed in three independent experiments on separate days. For endpoint analyses, spheroids from the three experiments were pooled (L3.6: n = 30 per condition; PANC-1: n = 15 per condition) and compared by unpaired t-test with Welch’s correction, as were the Ki-67 indices. Because spheroids were pooled across experiments, these endpoint p-values should be interpreted as exploratory. Migration kinetics were analyzed by repeated-measures two-way ANOVA (genotype × time; three spheroids per condition). Sample sizes for each panel are given in the figure legends.

Analyses used GraphPad Prism (version 10.6.1), R (version 4.4.2), or SAS (version 9.4; survival cut-points).

## Results

### *KRT17* marks a malignant epithelial population enriched for EMT- and migration-associated transcriptional programs

We first used scRNA-seq to identify the transcriptional programs associated with *KRT17* expression in malignant epithelial cells, compiled from two independent PDAC datasets comprising a total of 42 patients. Following quality control, integration, and malignant-cell identification, 77,155 malignant epithelial cells were retained for downstream analysis (Figure 1A-B). Within the malignant compartment, *KRT17* expression was distributed along a continuous gradient rather than a discrete binary population (Figure 1C), suggesting that *KRT17* may represent a biological cell state rather than a separate cellular lineage.

To identify biological programs associated with *KRT17* expression, malignant epithelial cells were stratified into *KRT17*-positive and *KRT17*-negative groups and subjected to differential expression and pathway-enrichment analyses. Hallmark gene-set enrichment analysis identified EMT as the most strongly represented pathway among *KRT17*-positive cells, exhibiting the highest normalized enrichment score of all cancer-related Hallmark pathways examined (NES = 3.06; Figure 1D-E) and the highest gene ratio (Supplementary Figure S1C). Additional pathways associated with *KRT17-*positive cells included MYC Targets V1 and TNFα signaling via NF-κB. Similar findings were obtained using independent Gene Ontology gene sets related to EMT and positive regulation of epithelial-cell migration (Supplementary Figure S1A-B).

To identify EMT/migration-associated markers linked with the *KRT17*-positive state, differentially expressed genes were intersected with a curated EMT/migration gene set comprising Hallmark EMT together with EMT- and migration-related Gene Ontology annotations. This analysis revealed substantial enrichment of EMT/migration-associated genes within *KRT17*-positive cells (265 genes) compared with *KRT17*-negative cells (68 genes) (Figure 1F). From this group, six biologically relevant markers were selected for downstream validation, namely, ANXA1, S100A4, GPC1, LAMC2, WWTR1, and LAMA3 (Figure 1G-H). Each marker exhibited increased expression prevalence within *KRT17*-positive cells and demonstrated positive single-cell correlation with *KRT17* expression (Spearman ρ = 0.16-0.48; Supplementary Figure S1D-J). The PCA plots showing their distribution in the malignant epithelial cells are in Supplementary Figure S1K-P.

Collectively, these analyses indicate that *KRT17* marks a malignant epithelial population characterized by EMT- and migration-associated transcriptional programs and identify six *KRT17*-associated markers for downstream validation. The selection was guided not only by their positive correlations with K17, but also by published literature on their roles in EMT and the availability of antibodies compatible with multiplex immunofluorescence.

### K17-positive PDAC tumors exhibit an EMT-associated proteomic phenotype

To determine whether the biological programs associated with K17 expression were also evident at the protein level, we analyzed proteomic data from 140 treatment-naive PDAC patients profiled by the CPTAC (Table 1). Tumors were stratified according to K17 protein abundance and subjected to differential protein abundance and pathway-enrichment analyses.

Consistent with the transcriptomic findings, Hallmark pathway analysis identified EMT as the most strongly represented biological program among K17-positive tumors, ranking first among all significantly enriched pathways (Figure 2A). The Hallmark EMT protein signature demonstrated highly significant enrichment within K17-positive tumors (NES = 3.041, FDR = 6.21 × 10⁻²⁸; Figure 2B), providing independent support for the association between K17 expression and an EMT-associated phenotype.

**Figure 2.**
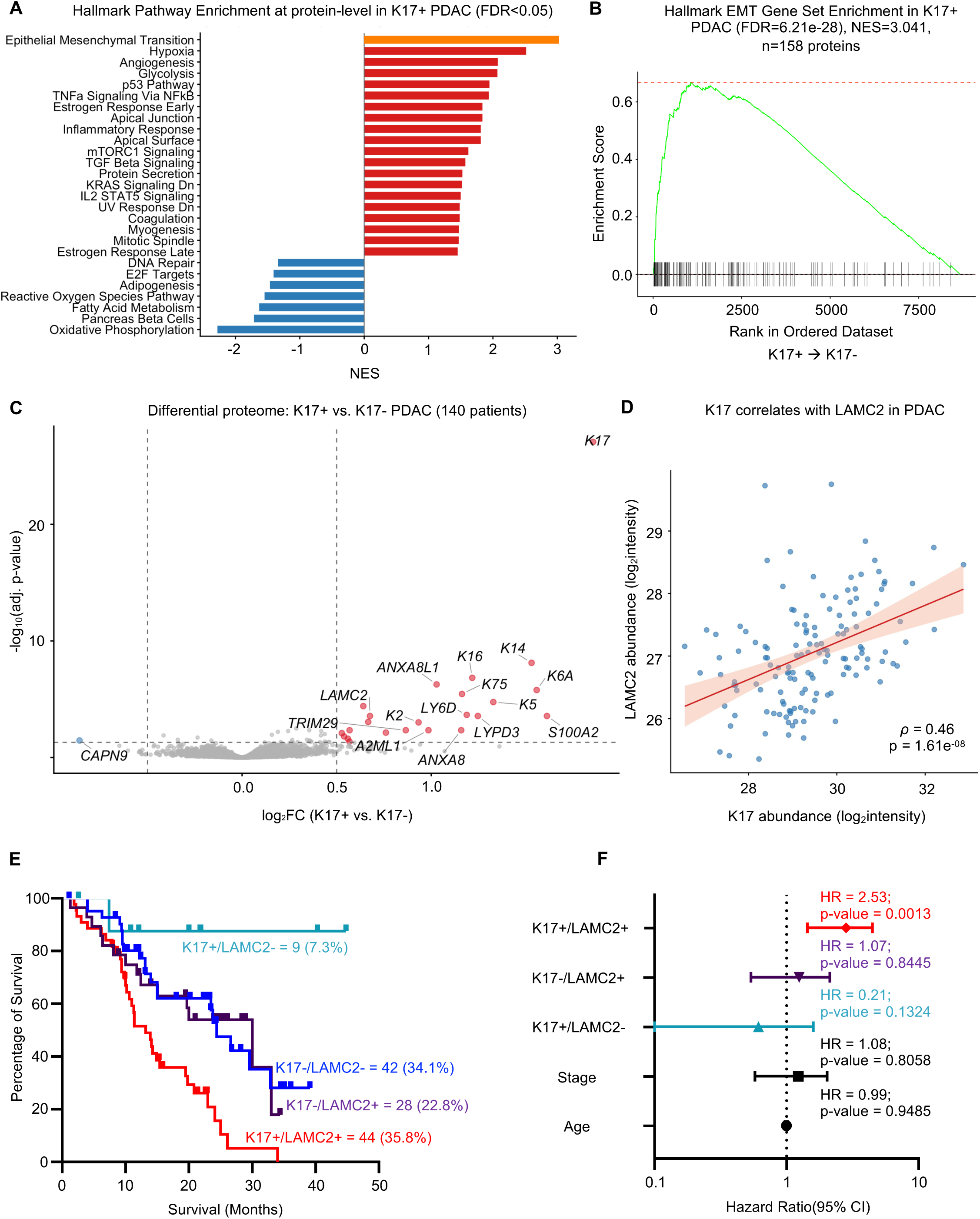
CPTAC proteomic analyses identify LAMC2 as the strongest protein-level correlate of K17 among the six scRNA-seq-nominated markers and associate K17/LAMC2 co-expression with adverse survival. (A) Enrichment of 27 cancer-related hallmark pathways in K17-positive vs. K17-negative tumors (140 patients; FDR < 0.05); EMT ranks first. (B) GSEA enrichment plot for the Hallmark EMT gene set (NES = 3.041; FDR = 6.21 × 10⁻²⁸; n = 158 proteins). (C) Volcano plot of differential protein abundance, K17-positive vs. K17-negative tumors; LAMC2 (log2FC = 0.676; adjusted p = 0.0002) and additional keratins/basal/EMT-associated markers are labeled among the top differentially abundant proteins. (D) K17 protein abundance correlates with LAMC2 abundance (n = 140 patients; Spearman ρ = 0.46; p = 1.61 × 10⁻⁸). (E) Kaplan-Meier overall survival by four-group K17/LAMC2 co-expression status (n = 123 patients); the K17-positive/LAMC2-positive group (n = 44 patients) shows the worst survival. (F) Forest plot of a multivariable Cox proportional hazards model for overall survival, including K17/LAMC2 co-expression groups (reference: K17-negative/LAMC2-negative) adjusted for pathologic stage and age; K17-positive/LAMC2-positive remained the only significant independent predictor of worse survival (HR = 2.53; p = 0.0013).

Differential protein abundance analysis identified LAMC2 among the significantly upregulated proteins in K17-positive tumors (log2FC = 0.676; adjusted p = 0.0002; Figure 2C). We next examined the relationship between K17 abundance and each of the six markers emerging from the single-cell analysis. Whereas GPC1 and LAMA3 demonstrated modest positive correlations with K17 abundance, ANXA1, S100A4, and WWTR1 did not show significant protein-level correlations (Supplementary Figure S2A-E). LAMC2 showed the strongest protein-level association with K17 among the six shortlisted markers (Spearman ρ = 0.46, p = 1.61 × 10⁻⁸; Figure 2D).

Together, these data support an association between K17 expression and an EMT-associated proteomic phenotype and identify LAMC2 as the strongest protein-level correlate of K17 among the six candidates.

### Combined K17/LAMC2 expression is associated with adverse overall survival

To explore the potential clinical relevance of the K17-LAMC2 relationship, survival analyses were performed using the subset of CPTAC patients with available survival data (n = 123). Patients were stratified by combined K17 and LAMC2 status, and survival was analyzed with Kaplan-Meier and multivariable Cox proportional hazards models.

Patients with K17-positive/LAMC2-positive tumors exhibited significantly worse overall survival than patients with K17-negative/LAMC2-negative tumors (adjusted HR = 2.53, p = 0.0013; Figure 2E-F). Neither K17-positive/LAMC2-negative nor K17-negative/LAMC2-positive groups differed significantly from the reference group. Individually, both K17 positivity (HR = 1.93, p = 0.009) and LAMC2 positivity (HR = 2.01, p = 0.004) were associated with worse survival in univariate analyses (Supplementary Figure S2F-H).

Given the discovery-cohort nature of these analyses and the relatively small size of some subgroup comparisons, these findings should be regarded as exploratory and will require validation in larger independent patient cohorts. Nevertheless, the results suggest that combined K17/LAMC2 expression may identify a particularly high-risk subset of PDAC patients.

### K17 is enriched in poorly differentiated PDAC regions and co-expressed with LAMC2 and ANXA1

To validate our transcriptomic and proteomic findings at spatial, single-cell resolution, we performed multiplex immunofluorescence imaging of PDAC TMA using the MACSima™ platform, staining for PanCK, K17, ANXA1, S100A4, GPC1, LAMC2, WWTR1, and LAMA3 across 69 tissue cores from 63 patients (tumor cell segmentation parameters summarized in Supplementary Table S3). Representative images from ten ROIs, drawn from ten distinct cores and ordered from lowest to highest intratumoral grade III content, illustrated the substantial intratumoral morphologic heterogeneity of PDAC. ROIs at the lower end of this progression were predominantly composed of grade I and grade II glandular architecture (Figure 3A), transitioning through intermediate ROIs containing varying admixtures of grade II and grade III morphology (Figure 3B-I), to an ROI composed predominantly of grade III, poorly differentiated tumor cell clusters (Figure 3J). Across this progression, K17 was qualitatively observed within grade II and grade III tumor components, but was largely absent from grade I-like, well-differentiated glandular structures.

**Figure 3.**
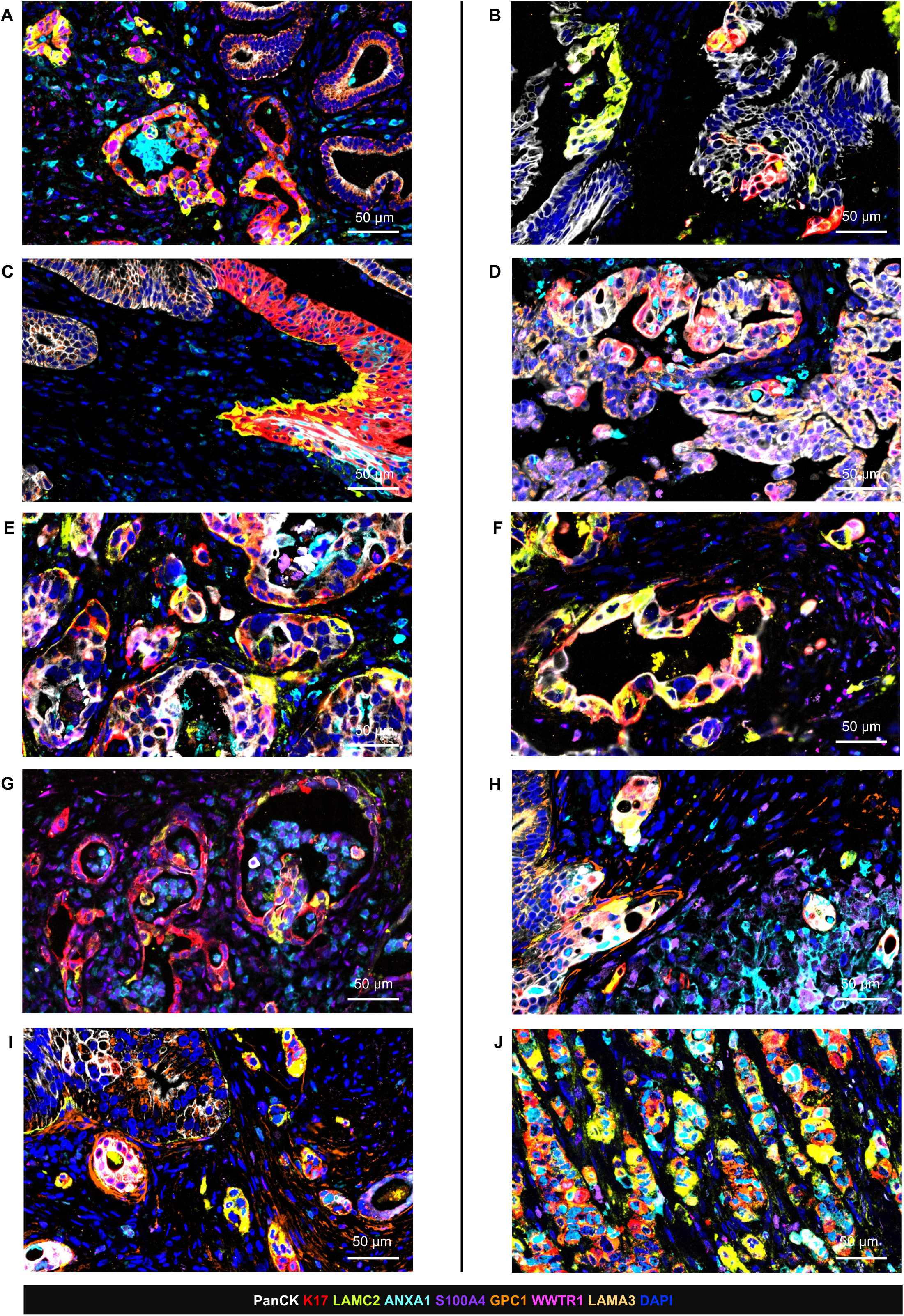
Multiplex spatial imaging (MACSima) shows K17 enrichment in moderately to poorly differentiated regions of PDAC. (A-J) Representative multiplex images from ten regions of interest (ROIs), drawn from ten distinct tissue cores and ordered from lowest to highest proportion of poorly differentiated content, showing PanCK (white), K17 (red), and the six EMT/migration-associated markers: LAMC2 (green), ANXA1 (teal), S100A4 (purple), GPC1 (orange), WWTR1 (pink), LAMA3 (beige) with DAPI counterstain (blue). The progression spans predominantly glandular architecture (A) through ROIs with varying admixtures of morphology (B-I) to an ROI composed predominantly of poorly differentiated tumor cell clusters lacking glandular architecture (J). K17 is primarily observed within moderately to poorly differentiated tumor components across this progression, and along with its co-expression with LAMC2, it gives a yellow color. Scale bars, 50 μm.

To quantify the relationship between K17 and the candidate markers at single-cell resolution, MACSima intensity data for all markers other than PanCK, restricted to tumor cells, were clustered with the Leiden algorithm, yielding 38 clusters, as visualized by a UMAP embedding (Figure 4A; N = 66 of 69 total cores with complete-panel staining, 410,605 tumor cells). K17 positivity was concentrated within a discrete subset of clusters rather than distributed uniformly across the UMAP (Figure 4B). Ordering clusters by K17 positivity and examining marker enrichment across them (Figure 4C, left) revealed that ANXA1 and LAMC2 tracked most closely with K17, supported by cluster-level Spearman correlation with K17: ANXA1 (ρ = 0.68) and LAMC2 (ρ = 0.49) showed the strongest positive correlations, followed by WWTR1 (ρ = 0.24), S100A4 and LAMA3 (both ρ = 0.15). GPC1 showed a weak inverse correlation (ρ = −0.20), being enriched in K17-negative rather than K17-positive clusters, a reversal from its nomination as a positively K17-associated gene at the transcript level (Figure 4C, right). Representative multiplex images confirmed spatial co-expression of K17 with both LAMC2 and ANXA1 at the single-cell level (Figure 4D, E). Their single-channel images along with PanCK are present in Supplementary Figure S4K-N. Quantitatively, the percentage of LAMC2-positive cells was significantly greater within K17-positive relative to K17-negative tumor cells (N = 67 cores, p < 0.0001; Figure 4F), and the same was true for ANXA1 (N = 69 cores, p < 0.0001; Figure 4G). Among the remaining candidates, GPC1 showed a significant difference in co-expression with K17 (N = 69 cores, p < 0.0001; Supplementary Figure S4H), consistent in direction with its inverse cluster-level correlation. S100A4 (N = 68 cores; Supplementary Figure S4G), WWTR1 (N = 69 cores; Supplementary Figure S4I), and LAMA3 (N = 69 cores; Supplementary Figure S4J) showed no significant co-expression with K17. The spatial distribution of each marker relative to K17-positive clusters was additionally shown by individual UMAP feature plots (Supplementary Figure S4A-F).

**Figure 4.**
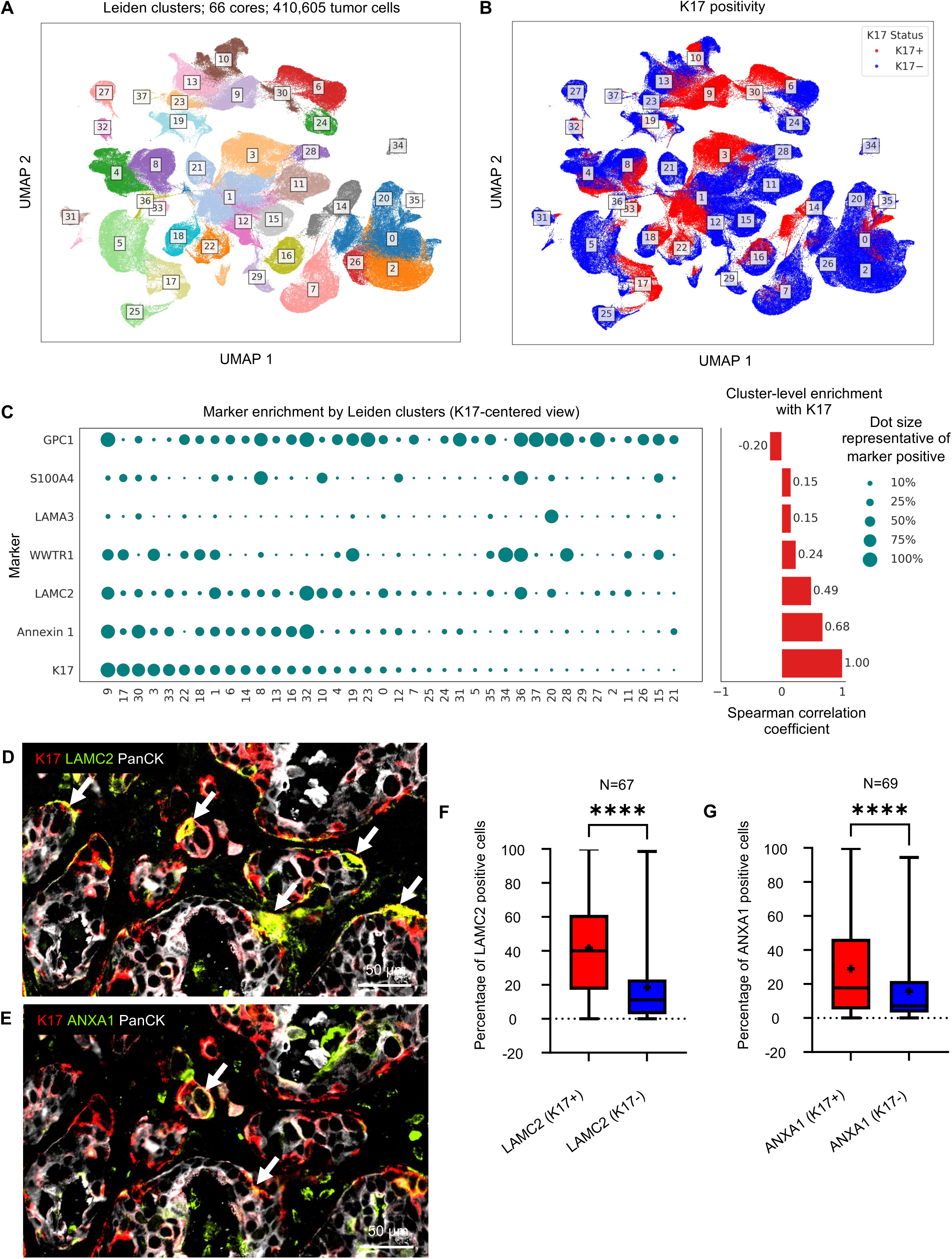
Single-cell-resolution MACSima analysis identifies LAMC2 and ANXA1 as the markers most strongly co-expressed with K17. (A) Leiden clustering of MACSima single-cell intensity data of all six EMT-associated markers and K17 (excluding PanCK), restricted to tumor cells, visualized by UMAP (N = 66 of 69 total cores with complete-panel staining; 410,605 tumor cells; 38 clusters). (B) K17 positivity overlaid on the same UMAP, with K17-positive cells concentrated within a discrete subset of clusters. (C) Marker enrichment across clusters ordered by K17 positivity (left) and cluster-level Spearman correlation of each marker with K17 (right): ANXA1 ρ = 0.68, LAMC2 ρ = 0.49, WWTR1 ρ = 0.24, S100A4 ρ = 0.15, LAMA3 ρ = 0.15, GPC1 ρ = −0.20. (D) Representative multiplex image showing K17 (red), LAMC2 (green), and PanCK (white) merge; arrows indicate co-expressing cells. (E) Representative multiplex image showing K17 (red), ANXA1 (green), and PanCK (white) merge; arrows indicate co-expressing cells. (F) Percentage of LAMC2-positive cells in K17-positive vs. K17-negative tumor cells (N = 67 cores, ****p < 0.0001). (G) Percentage of ANXA1-positive cells in K17-positive vs. K17-negative tumor cells (N = 69 cores, ****p < 0.0001). Boxes show the median and interquartile range, whiskers show the range, and “+” marks the mean. Wilcoxon matched-pairs signed-rank test. Scale bars, 50 μm.

In summary, K17 localized to poorly differentiated regions in representative images and, at single-cell resolution, was most strongly associated with LAMC2 and ANXA1.

### K17/LAMC2 co-expression increases with intratumoral dedifferentiation

Dual-plex immunohistochemistry for K17 and LAMC2 was performed on whole-slide sections from an overlapping cohort of 60 PDAC cases. Tumor area within each case was stratified by morphologic intratumoral grade (as defined above), independent of the whole-tumor World Health Organization (WHO) grade assigned during case selection (Figure 5A-C) and further subdivided into four categories based on combined K17/LAMC2 IHC status: K17-positive/LAMC2-positive, K17-positive/LAMC2-negative, K17-negative/LAMC2-positive, and K17-negative/LAMC2-negative (Figure 5D-F). Most tumors contained a mixture of intratumoral grades rather than a single uniform grade (Figure 5G). Within each intratumoral grade compartment, the K17-positive/LAMC2-positive category occupied a progressively greater proportion of tumor area moving from grade I to grade III, while the K17-negative/LAMC2-negative category showed the inverse pattern, occupying proportionally less area with worsening grade (Figure 5H-J). This grade-dependent shift was significant for all four K17/LAMC2 categories by Friedman test (K17-positive/LAMC2-positive: χ² = 93.11, p = 6.04 × 10⁻²¹; K17-negative/LAMC2-positive: χ² = 72.45, p = 1.85 × 10⁻¹⁶; K17-positive/LAMC2-negative: χ² = 55.37, p = 9.48 × 10⁻¹³; K17-negative/LAMC2-negative: χ² = 24.68, p = 4.37 × 10⁻⁶), and post hoc pairwise Wilcoxon signed-rank testing confirmed that every grade-to-grade comparison across all four categories remained significant after Benjamini-Hochberg correction across the full set of 12 comparisons (all adjusted p < 0.05, including the full-range grade I versus grade III comparison for K17-positive/LAMC2-positive: adjusted p = 2.22 × 10⁻¹⁰). Because grade I, II, and III measurements were drawn from the same 60 cases, these findings describe intratumoral spatial heterogeneity: individual tumors shift toward K17-positive/LAMC2-positive composition specifically within their less-differentiated regions, and this pattern was evident at whole-slide scale. Single-plex IHC of K17 and LAMC2 on adjacent serial sections further confirmed direct spatial overlap between the two markers within the same tumor architecture (Supplementary Figure S5).

**Figure 5.**
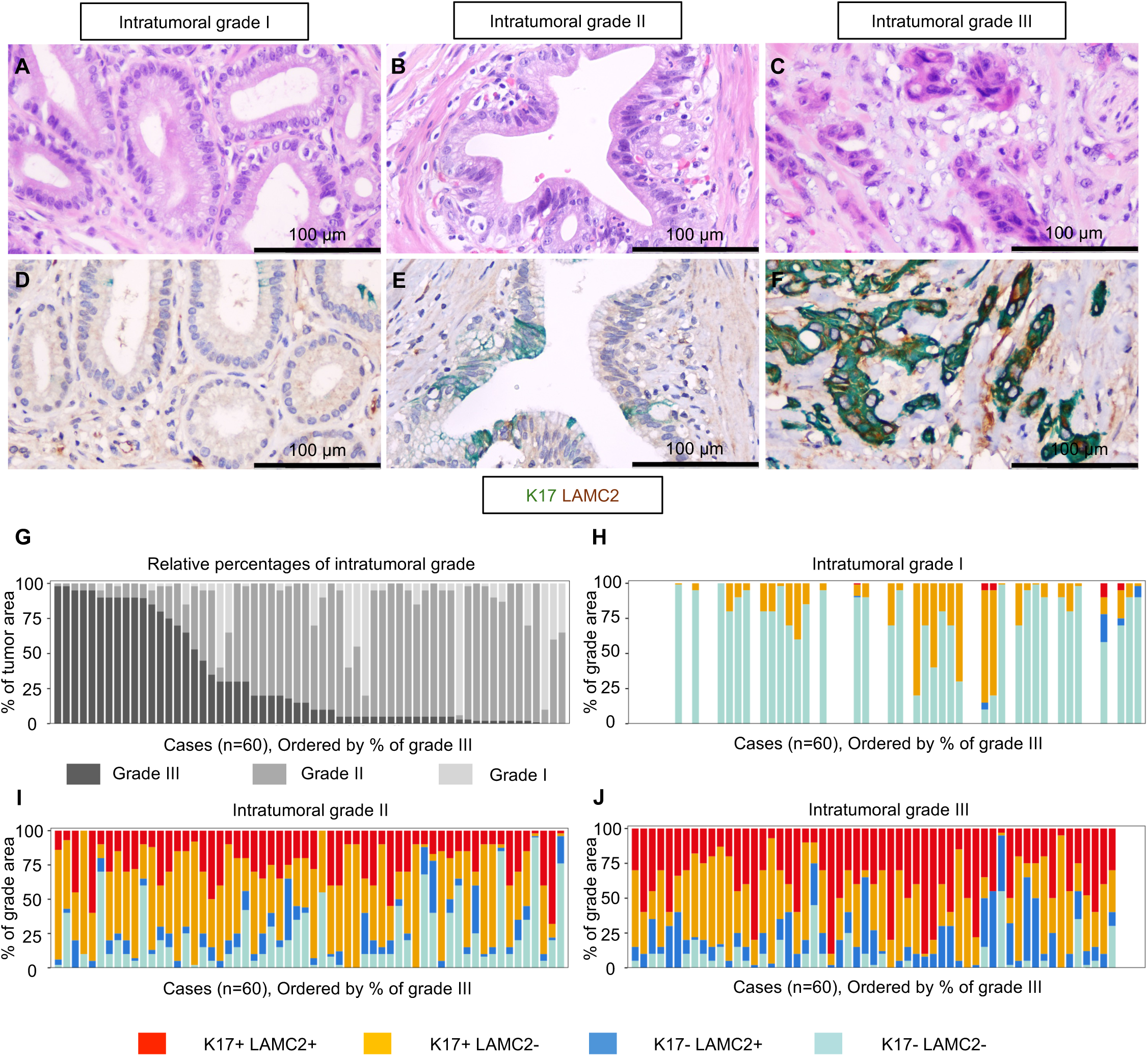
K17/LAMC2 co-expression tracks with intratumoral histologic grade. (A-C) Representative H&E images of intratumoral grade I, II, and III morphology. (D-F) Corresponding dual-plex IHC for K17 (green) and LAMC2 (brown) in an adjacent serial section. (G) Relative percentage of intratumoral grade (I/II/III) per case (n = 60), ordered by percent intratumoral grade III area. (H-J) Percentage of grade area occupied by each of four K17/LAMC2 categories within grade I (H), grade II (I), and grade III (J), showing progressive enrichment of the K17-positive/LAMC2-positive category and progressive loss of the K17-negative/LAMC2-negative category with worsening grade (Friedman test with Benjamini-Hochberg-corrected pairwise comparisons; all grade-to-grade comparisons significant, adjusted p < 0.05). Scale bars, 100 μm.

These findings indicate that K17/LAMC2 co-expression is associated with tumor dedifferentiation and becomes increasingly prevalent within poorly differentiated PDAC regions.

### K17 promotes migratory outgrowth in three-dimensional PDAC spheroid models

Having established a consistent association between K17 expression and an EMT-associated phenotype across transcriptomic, proteomic, and spatial analyses, we next sought to determine whether K17 influences tumor-cell migratory behavior in three-dimensional collagen I spheroid migration assays.

L3.6 cells were used to generate isogenic loss-of-function (K17-knockout or KO) and gain-of-function (K17-overexpression or K17) models. PANC-1 cells, which do not express endogenous K17, were used as an independent gain-of-function model. In L3.6 cells, K17 knockout reduced total spheroid area, reflecting both the spheroid core and its migratory outgrowth combined, over time relative to wild-type (WT) controls (two-way ANOVA, p = 0.0478; Figure 6A-B), while K17 overexpression increased total spheroid area relative to empty vector (EV) controls (two-way ANOVA, p = 0.0026; Figure 6C-D), an effect that was recapitulated in PANC-1 cells (two-way ANOVA, p = 0.0004; Figure 6E-F). To isolate migratory behavior specifically from overall growth, a normalized migration index was calculated at the endpoint of each experiment (96 hours for L3.6, 48 hours for PANC-1) (see Methods). KO L3.6 spheroids showed a significantly reduced migration index relative to WT (p < 0.0001; Figure 6G-H), while K17-overexpressing L3.6 and PANC-1 spheroids each showed a significantly increased migration index relative to their EV controls (p < 0.0001; Figure 6I-L). Confocal imaging of whole-mount spheroids showed K17 expression, predominantly at the periphery, in WT L3.6 and in K17-overexpressing L3.6 and PANC-1 spheroids (Supplementary Figure S6A-C). However, immunohistochemistry of FFPE sections of spheroids from the same experiment showed K17 staining throughout the full depth of the K17 spheroid, including the core (Supplementary Figure S6D, E), suggesting that this peripherally restricted confocal signal reflects limited antibody and light penetration into the intact spheroid rather than true spatial restriction of K17 expression. To explore whether proliferation might contribute to these differences, Ki-67 labeling was assessed in parallel spheroids (Supplementary Figure S6F-K). Although KO spheroids demonstrated reduced migratory outgrowth, they exhibited increased Ki-67 labeling relative to WT controls (p = 0.0098). In contrast, Ki-67 labeling did not differ significantly between gain-of-function and control models (p = 0.2868 for L3.6 and 0.2912 for PANC-1). These observations suggest that differences in proliferation alone are unlikely to fully account for the migratory phenotype associated with K17 expression. These results support the conclusion that K17 is not merely a marker of the aggressive phenotype but also contributes to the migratory behavior in these PDAC models.

**Figure 6.**
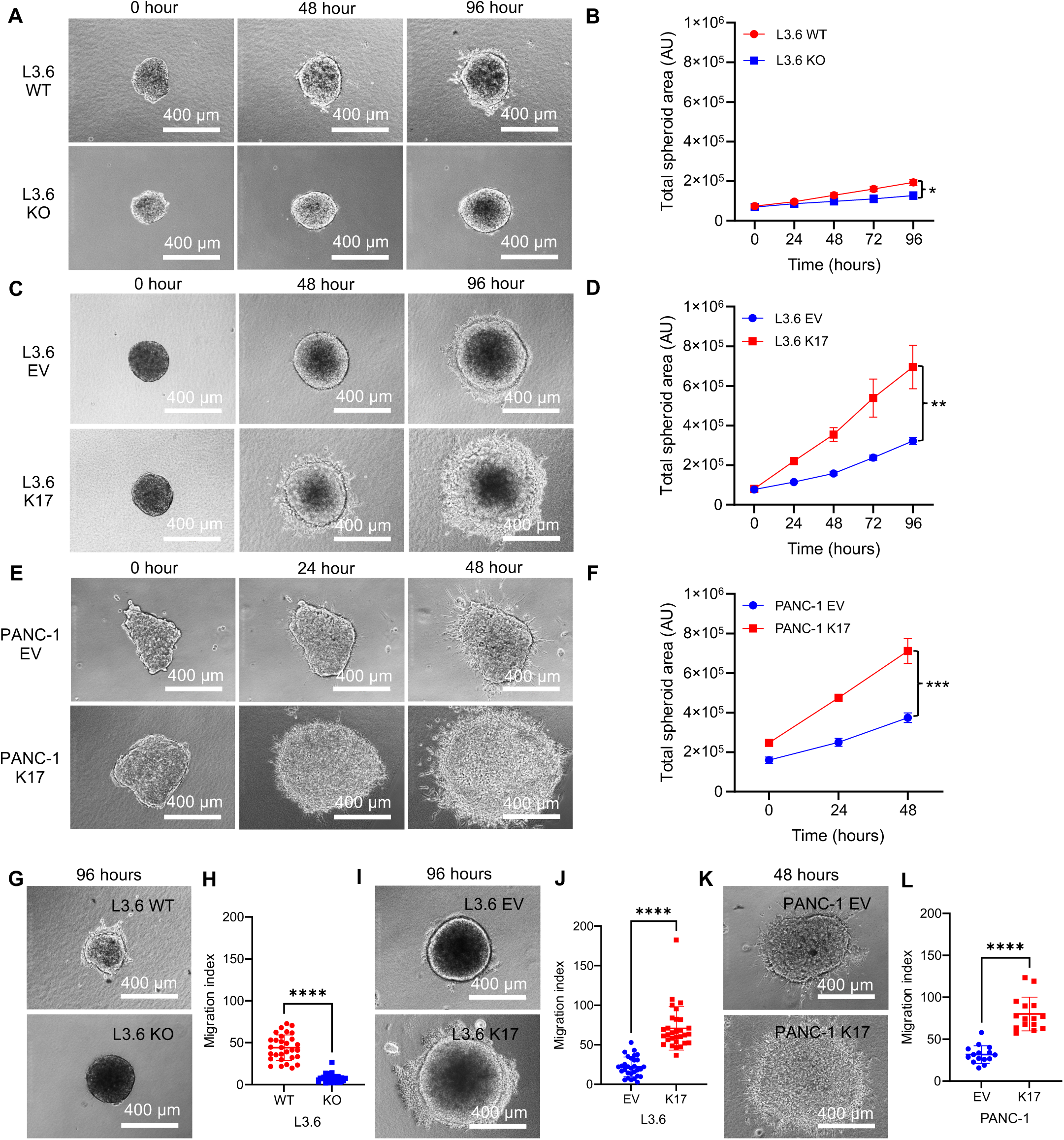
K17 promotes migratory outgrowth in three-dimensional PDAC spheroids. Representative spheroids were embedded in collagen I and monitored by phase-contrast microscopy over time (96 hours for L3.6 and 48 hours for PANC-1). (A, B) L3.6 wild-type (WT) versus K17-knockout (KO) spheroids: representative images and quantification of total spheroid area over time (two-way ANOVA, n = 3, p = 0.0478). (C, D) L3.6 empty-vector (EV) versus K17-overexpressing (K17) spheroids: representative images and quantification of total spheroid area over time (two-way ANOVA, n = 3, p = 0.0026). (E, F) PANC-1 EV versus K17 spheroids: representative images and quantification of total spheroid area over time (two-way ANOVA, n = 3, p = 0.0004). (G, H) Migration index of L3.6 WT versus KO spheroids at 96 hours. (I, J) Migration index of L3.6 EV versus K17 spheroids at 96 hours. (K, L) Migration index of PANC-1 EV versus K17 spheroids at 48 hours. Endpoint migration indices were derived from three independent biological experiments, with individual spheroids pooled and analyzed (L3.6: *n* = 30 spheroids per condition; PANC-1: *n* = 15 spheroids per condition). Statistical comparisons of endpoint migration indices were performed using an unpaired t-test with Welch’s correction (****p < 0.0001). Data are presented as mean ± SD. Scale bars, 400 μm.

## Discussion

K17 is a well-established marker of the aggressive basal-like subtype of PDAC and an independent predictor of poor survival.^4^ Yet the biological basis for this association has remained incompletely understood. By integrating single-cell transcriptomics, proteomics, multiplex spatial imaging, histologic analysis, and functional experimentation, we show that K17 expression is associated with EMT-related transcriptional and protein-expression programs across independent datasets and analytical platforms, localized to morphologically less differentiated regions within tumors, and functionally promoted tumor-cell migration in three-dimensional PDAC models. Together, these findings provide a biological framework that links K17 expression to aggressive tumor behavior and may help explain its longstanding association with adverse clinical outcome.

One of the principal findings of this study is that K17-positive PDAC cells and tumors are enriched for EMT-associated programs at both the transcript and protein levels. EMT has been implicated in tumor dissemination, therapeutic resistance, and disease progression in PDAC^2,3^ and is a widely recognized program in carcinoma progression, although its requirement for metastasis in PDAC remains debated.^33–36^ Our findings place K17 within this biologically important context and suggest that K17 expression identifies a subset of tumor cells enriched for migratory and invasive properties. Rather than representing a general marker of aggressive disease, K17 appears to define a specific tumor-cell state characterized by molecular features commonly associated with EMT and tumor-cell motility.

Consistent with contemporary models of tumor plasticity, our findings are more compatible with an EMT-associated or partial-EMT state than with complete mesenchymal conversion.^37–40^ K17-positive cells were identified within the PanCK-positive epithelial tumor compartment while showing enrichment for genes and proteins associated with migration, extracellular-matrix interaction, and invasion, a pattern consistent with a partial rather than complete EMT, although canonical epithelial and mesenchymal markers were not directly assessed. This observation is consistent with growing evidence that partial EMT states may be particularly relevant to tumor progression because they preserve epithelial features that facilitate survival and collective migration while simultaneously acquiring characteristics associated with invasion and metastatic competence. This pattern is therefore consistent with models in which PDAC progression is shaped by dynamic, plastic intermediate cell states rather than binary epithelial-to-mesenchymal transitions.

Independent of our own data, each of the six candidate markers nominated by the transcriptomic screen has prior literature support for a role in EMT or tumor invasion, providing biological plausibility for their selection. ANXA1 promotes migration and invasion in pancreatic cancer cell lines, both intracellularly and as a secreted extracellular form.^41^ S100A4 promotes PDAC cell motility and invasion and has been linked to Src/FAK signaling.^42^ GPC1 knockdown reverses EMT marker expression and reduces AKT/GSK/β-catenin signaling in another gastrointestinal adenocarcinoma model.^43^ WWTR1 (TAZ), a transcriptional effector of the Hippo pathway, confers EMT-associated, stem-cell-like properties in breast cancer and independently drives migration, invasion, and anchorage-independent growth.^44,45^ Lastly, LAMA3, a subunit of laminin-332, has been associated with EMT in cholangiocarcinoma cells.^46^ LAMC2 has been mechanistically linked to PDAC tumorigenesis through EGFR/ERK1/2/AKT/mTOR signaling.^47^ It encodes the γ2 chain of laminin-332, whose proteolytic cleavage generates a pro-migratory signal^48^, and has been implicated in EMT, migration, invasion, and therapy resistance in PDAC and other cancers.^47,49–52^ Among these candidates, however, LAMC2 emerged as the most reproducible correlate of K17. Whereas several markers demonstrated associations within individual datasets or analytical platforms, LAMC2 remained linked to K17 throughout the entire validation pipeline, including transcriptomic analyses, protein-level correlation within CPTAC data, single-cell spatial co-expression by MACSima imaging, and whole-slide dual-plex IHC. This reproducibility across orthogonal approaches suggests that LAMC2 is not simply another EMT-associated marker but rather a robust companion marker of the K17-associated phenotype. ANXA1 showed a strong cluster-level spatial association with K17 (ρ = 0.68), but it did not do so at the protein level in CPTAC and was not evaluated for prognostic association. Among the other candidates, WWTR1, S100A4 and LAMA3 showed weak, non-significant associations with K17 on MACSima, and GPC1 was enriched in K17-negative tumor cells.

The relationship between K17 expression and tumor morphology represents another important finding of this study. The EMT-associated program was not uniformly distributed throughout tumors but instead concentrated within specific histologic regions. In representative MACSima regions of interest ordered by increasing intratumoral grade III content, K17 was largely absent from well-formed, grade I-like glandular structures and progressively enriched within grade II and grade III components. Whole-slide dual-plex IHC of 60 tumors quantitatively confirmed this pattern, with K17-positive/LAMC2-positive tumor area occupying a progressively larger proportion of each grade compartment from grade I to grade III (Friedman χ² = 93.11, p = 6.04 × 10⁻²¹). The concordance between single-cell spatial imaging and whole-slide histologic analysis argues against a platform-specific artifact and supports the conclusion that K17 expression is linked to regional loss of glandular differentiation within individual tumors.

These findings were observed within individual tumors rather than solely across different tumors, indicating that K17 is linked to intratumoral morphologic heterogeneity. Thus, K17 may not simply serve as a surrogate marker of high-grade PDAC but instead identify a biologically distinct population of tumor cells that associates with dedifferentiation and aggressive tumor behavior. These findings also emphasize that molecular and morphologic progression are closely linked processes in PDAC and suggest that K17-positive cells occupy a recognizable histologic niche within the tumor ecosystem.

Combined K17/LAMC2 positivity identified the CPTAC subset with the poorest overall survival (adjusted HR = 2.53 versus double-negative tumors, p = 0.0013; n = 123), and this double-positive population expanded within poorly differentiated regions on dual-plex IHC. These analyses remain exploratory because they relied on protein-specific optimal cut-points, involved relatively small subgroups, and were restricted to a single cohort. We therefore view LAMC2 primarily as a reproducible companion marker of the K17-associated phenotype.

Our findings are also consistent with the emerging concept that PDAC aggressiveness is shaped by dynamic cell states in addition to static molecular subtypes. K17 expression was distributed along a continuum rather than defining a discrete population in the single-cell data, suggesting that K17 reflects a graded, potentially plastic transcriptional state. This observation aligns with evidence that classical and basal-like programs coexist within individual tumors and along a continuum^12–14^ and with reports in other epithelial cancers that specialized, plastic tumor-cell states drive aggressive behavior.^53^ As a working model, K17 identifies cells positioned toward the migratory, dedifferentiated end of this phenotypic spectrum.

Whether K17 simply identifies this aggressive cellular state or actively contributes to its biological properties remains one of the most important unanswered questions surrounding K17 biology. Using three-dimensional collagen I-embedded spheroids, which better model matrix engagement and collective cell migration than conventional monolayer systems, we found that K17 knockout reduced whereas K17 overexpression increased migratory outgrowth. Ki-67 staining did not identify corresponding differences in proliferation that could account for these observations (Figure 6; Supplementary Figure S6). The bidirectional effects observed in gain- and loss-of-function models support a functional contribution of K17 to migratory behavior and argue against K17 serving solely as a passive biomarker of aggressive disease. While these experiments do not establish the downstream signaling mechanisms involved, they demonstrate that altering K17 expression alone is sufficient to modify tumor-cell migration in a biologically relevant three-dimensional environment.

Although this study establishes a role for K17 in migration *in vitro*, the molecular mechanisms remain undefined. K17 has been linked to p27 nuclear export and degradation in cervical, breast and pancreatic cancer cells, to immune modulation of the PDAC microenvironment, to mitochondrial pyrimidine biosynthesis and chemoresistance in basal-like PDAC, and broader roles are reviewed by Baraks et al..^6–8,54^ Among other epithelial cancers, K17 induces EMT through AKT activation in esophageal squamous cell carcinoma and bladder cancer, and Wnt/β-catenin signaling has been implicated in K17-associated doxorubicin resistance in triple-negative breast cancer.^55–57^ Future studies will be needed to determine how K17 interacts with cytoskeletal, adhesion, and extracellular matrix signaling pathways that regulate tumor-cell movement. It remains to be tested whether AKT signaling, which LAMC2 has been reported to activate in pancreatic cancer cells, links K17 and LAMC2; our data do not address this question.^50^

Several limitations should be acknowledged. The analyses of human tissues remain largely correlative, and the MACSima and dual-plex IHC cohorts were overlapping and derived from a single institution. Canonical EMT regulators such as vimentin, ZEB1, SNAIL, and SLUG were not directly interrogated and were not included in the final MACSima panel. Functional validation was limited to two PDAC cell lines and was confined to in vitro experimentation. Additional studies in independent clinically annotated cohorts and complementary experimental systems will therefore be necessary to establish the generalizability and mechanistic basis of these observations. Mechanistic studies will be required to determine how K17 and LAMC2 are linked and whether K17 regulates EMT-associated signaling.

Despite these limitations, the consistency of findings across independent single-cell transcriptomic datasets, proteomic analyses, multiplex spatial imaging, histopathology, and functional experiments supports the conclusion that K17 identifies a biologically meaningful tumor-cell state in PDAC. Across these orthogonal approaches, K17 was repeatedly linked to EMT-associated programs, progressive loss of glandular differentiation, and enhanced migratory behavior. These observations offer a possible biological context for the longstanding association between K17 expression, poor differentiation, and adverse clinical outcome and support a model in which K17-positive cells represent an aggressive component of the tumor ecosystem.

## Conclusion

This study identifies K17 as a marker of a poorly differentiated, EMT-associated migratory cell state in PDAC. Across independent single-cell, proteomic, spatial, histologic, and functional analyses, K17 consistently tracked with EMT-related programs, localized to higher-grade regions within tumors, and promoted migration in three-dimensional experimental models. LAMC2 emerged as the most reproducible companion marker of this phenotype across multiple analytical platforms. Collectively, these findings provide a biological explanation for the adverse prognostic significance of K17 and support a model in which K17-positive cells represent a dynamic, aggressive tumor-cell population that contributes to PDAC progression.

## Data availability

The scRNA-seq datasets analyzed in this study are publicly available and were originally published by Peng et al. (2019, Cell Research) and Werba et al. (2023, Nature Communications). Raw sequencing data from Peng et al. are available in the Genome Sequence Archive (GSA) under accession CRA001160, and data from Werba et al. are available in the Gene Expression Omnibus (GEO) under accession GSE205013. Proteomic data for the CPTAC PDAC cohort are publicly available through the LinkedOmics data portal (linkedomics.org), corresponding to the dataset originally described by Cao et al. (2021, *Cell*). The MACSima multiplex immunofluorescence imaging data, dual-plex and single-plex immunohistochemistry images, 3D spheroid migration assay images, and all associated processed data (including single-cell intensity matrices, quantification outputs, and analysis code) generated during this study are available from the corresponding author upon reasonable request. All code used for computational analyses, including scRNA-seq processing, differential expression analysis, gene set enrichment analysis, and survival analysis, is available from the corresponding author upon reasonable request.

## Authors’ contributions

**S. Sarkar**: Writing - original draft; Review & editing; Conceptualization; Investigation; Data curation; Formal analysis; Methodology; Visualization.

**M. Ghosh**: Review & editing; Conceptualization; Methodology; Formal analysis.

**L. Delgado-Coka**: Review & editing; Resources; Data curation; Methodology.

**S. Powers**: Review & editing; Data curation; Formal analysis; Visualization.

**R. Sathian**: Review & editing; Formal analysis; Visualization.

**R. Davuluri**: Formal analysis.

**X. Gong**: Review & editing; Formal analysis; Resources; Methodology.

**M. Mak**: Review & editing; Resources; Methodology.

**J. Villar**: Methodology.

**X. Qi**: Formal analysis.

**S. Babu**: Formal analysis.

**N. Marchenko**: Review & editing; Conceptualization; Supervision.

**K.R. Shroyer**: Review & editing; Conceptualization; Investigation; Supervision; Funding acquisition.

## Acknowledgements

The authors are grateful to members of the Shroyer laboratory for their thoughtful discussions and support throughout the course of this work. We thank Germana Giupponi for her contributions in gathering the clinical data for the PDAC cases used in this study, and Dr. Wei-Xing Zong (Rutgers University) for providing the L3.6 cell line. We also thank the Biobank at Stony Brook University Hospital for providing archival PDAC tissue specimens. Claude (Anthropic) and Copilot (Microsoft) were used to assist with language polishing; while only Claude (Anthropic) was used for code refinements during manuscript preparation and analysis. This work was supported by Academic Enrichment Funds of the Department of Pathology at the Renaissance School of Medicine, the Marvin Kuschner Endowed Professorship, and Pancreatic Cancer Action Network Translational Research Grant 18-65-SHRO (to K.R. Shroyer). We acknowledge support from NIH grant R35GM142875 (to M. Mak).

## Ethics approval and consent

The use of archival human tissue specimens was approved by the Stony Brook University Institutional Review Board (IRB ID: 94651_MODCR016), which waived the requirement for informed consent because the study was retrospective and used de-identified archival specimens. The study was conducted in accordance with the Declaration of Helsinki. The scRNA-seq and CPTAC proteomic analyses used publicly available, de-identified datasets and required no additional ethical approval.

## References

1 Siegel, R. L., Kratzer, T. B., Giaquinto, A. N., Sung, H. & Jemal, A. Cancer statistics, 2025. CA Cancer J Clin 75, 10–45 (2025). 10.3322/caac.21871

2 Rhim, A. D. et al. EMT and dissemination precede pancreatic tumor formation. Cell 148, 349–361 (2012). 10.1016/j.cell.2011.11.025

3 Arumugam, T. et al. Epithelial to mesenchymal transition contributes to drug resistance in pancreatic cancer. Cancer Res 69, 5820–5828 (2009). 10.1158/0008-5472.Can-08-2819

4 Roa-Peña, L. et al. Keratin 17 identifies the most lethal molecular subtype of pancreatic cancer. Sci Rep 9, 11239 (2019). 10.1038/s41598-019-47519-4

5 Escobar-Hoyos, L. F. et al. Keratin 17 in premalignant and malignant squamous lesions of the cervix: proteomic discovery and immunohistochemical validation as a diagnostic and prognostic biomarker. Mod Pathol 27, 621–630 (2014). 10.1038/modpathol.2013.166

6 Baraks, G. et al. Dissecting the Oncogenic Roles of Keratin 17 in the Hallmarks of Cancer. Cancer Res 82, 1159–1166 (2022). 10.1158/0008-5472.Can-21-2522

7 Delgado-Coka, L. et al. Keratin 17 modulates the immune topography of pancreatic cancer. J Transl Med 22, 443 (2024). 10.1186/s12967-024-05252-1

8 Pan, C. H. et al. Mitochondrial-Localized Keratin 17 Promotes Chemoresistance in Basal-like Pancreatic Cancer. Cancer Res 86, 2935–2950 (2026). 10.1158/0008-5472.Can-25-4534

9 Collisson, E. A. et al. Subtypes of pancreatic ductal adenocarcinoma and their differing responses to therapy. Nat Med 17, 500–503 (2011). 10.1038/nm.2344

10 Moffitt, R. A. et al. Virtual microdissection identifies distinct tumor- and stroma-specific subtypes of pancreatic ductal adenocarcinoma. Nat Genet 47, 1168–1178 (2015). 10.1038/ng.3398

11 Bailey, P. et al. Genomic analyses identify molecular subtypes of pancreatic cancer. Nature 531, 47–52 (2016). 10.1038/nature16965

12 Raghavan, S. et al. Microenvironment drives cell state, plasticity, and drug response in pancreatic cancer. Cell 184, 6119–6137.e26 (2021). 10.1016/j.cell.2021.11.017

13 Chan-Seng-Yue, M. et al. Transcription phenotypes of pancreatic cancer are driven by genomic events during tumor evolution. Nat Genet 52, 231–240 (2020). 10.1038/s41588-019-0566-9

14 Williams, H. L. et al. Spatially Resolved Single-Cell Assessment of Pancreatic Cancer Expression Subtypes Reveals Co-expressor Phenotypes and Extensive Intratumoral Heterogeneity. Cancer Res 83, 441–455 (2023). 10.1158/0008-5472.Can-22-3050

15 Peng, J. et al. Single-cell RNA-seq highlights intra-tumoral heterogeneity and malignant progression in pancreatic ductal adenocarcinoma. Cell Res 29, 725–738 (2019). 10.1038/s41422-019-0195-y

16 Werba, G. et al. Single-cell RNA sequencing reveals the effects of chemotherapy on human pancreatic adenocarcinoma and its tumor microenvironment. Nat Commun 14, 797 (2023). 10.1038/s41467-023-36296-4

17 Zheng, G. X. et al. Massively parallel digital transcriptional profiling of single cells. Nat Commun 8, 14049 (2017). 10.1038/ncomms14049

18 Fleming, S. J. et al. Unsupervised removal of systematic background noise from droplet-based single-cell experiments using CellBender. Nat Methods 20, 1323–1335 (2023). 10.1038/s41592-023-01943-7

19 Wolf, F. A., Angerer, P. & Theis, F. J. SCANPY: large-scale single-cell gene expression data analysis. Genome Biol 19, 15 (2018). 10.1186/s13059-017-1382-0

20 Korsunsky, I. et al. Fast, sensitive and accurate integration of single-cell data with Harmony. Nat Methods 16, 1289–1296 (2019). 10.1038/s41592-019-0619-0

21 Linderman, G. C. et al. Zero-preserving imputation of single-cell RNA-seq data. Nat Commun 13, 192 (2022). 10.1038/s41467-021-27729-z

22 Liberzon, A. et al. The Molecular Signatures Database (MSigDB) hallmark gene set collection. Cell Syst 1, 417–425 (2015). 10.1016/j.cels.2015.12.004

23 Cao, L. et al. Proteogenomic characterization of pancreatic ductal adenocarcinoma. Cell 184, 5031–5052.e5026 (2021). 10.1016/j.cell.2021.08.023

24 Ritchie, M. E. et al. limma powers differential expression analyses for RNA-sequencing and microarray studies. Nucleic Acids Res 43, e47 (2015). 10.1093/nar/gkv007

25 Amin, M. B. et al. The Eighth Edition AJCC Cancer Staging Manual: Continuing to build a bridge from a population-based to a more “personalized” approach to cancer staging. CA Cancer J Clin 67, 93–99 (2017). 10.3322/caac.21388

26 Chun, Y. S., Pawlik, T. M. & Vauthey, J. N. 8th Edition of the AJCC Cancer Staging Manual: Pancreas and Hepatobiliary Cancers. Ann Surg Oncol 25, 845–847 (2018). 10.1245/s10434-017-6025-x

27 Nagtegaal, I. D. et al. The 2019 WHO classification of tumours of the digestive system. Histopathology 76, 182–188 (2020). 10.1111/his.13975

28 Traag, V. A., Waltman, L. & van Eck, N. J. From Louvain to Leiden: guaranteeing well-connected communities. Sci Rep 9, 5233 (2019). 10.1038/s41598-019-41695-z

29 Cong, L. et al. Multiplex genome engineering using CRISPR/Cas systems. Science 339, 819–823 (2013). 10.1126/science.1231143

30 Ran, F. A. et al. Genome engineering using the CRISPR-Cas9 system. Nat Protoc 8, 2281–2308 (2013). 10.1038/nprot.2013.143

31 Ilina, O. et al. Cell-cell adhesion and 3D matrix confinement determine jamming transitions in breast cancer invasion. Nat Cell Biol 22, 1103–1115 (2020). 10.1038/s41556-020-0552-6

32 Schindelin, J. et al. Fiji: an open-source platform for biological-image analysis. Nat Methods 9, 676–682 (2012). 10.1038/nmeth.2019

33 Thiery, J. P., Acloque, H., Huang, R. Y. & Nieto, M. A. Epithelial-mesenchymal transitions in development and disease. Cell 139, 871–890 (2009). 10.1016/j.cell.2009.11.007

34 Nieto, M. A., Huang, R. Y., Jackson, R. A. & Thiery, J. P. EMT: 2016. Cell 166, 21–45 (2016). 10.1016/j.cell.2016.06.028

35 Zheng, X. et al. Epithelial-to-mesenchymal transition is dispensable for metastasis but induces chemoresistance in pancreatic cancer. Nature 527, 525–530 (2015). 10.1038/nature16064

36 Aiello, N. M. et al. Upholding a role for EMT in pancreatic cancer metastasis. Nature 547, E7–e8 (2017). 10.1038/nature22963

37 Puram, S. V. et al. Single-Cell Transcriptomic Analysis of Primary and Metastatic Tumor Ecosystems in Head and Neck Cancer. Cell 171, 1611–1624.e1624 (2017). 10.1016/j.cell.2017.10.044

38 Pastushenko, I. et al. Identification of the tumour transition states occurring during EMT. Nature 556, 463–468 (2018). 10.1038/s41586-018-0040-3

39 Aiello, N. M. et al. EMT Subtype Influences Epithelial Plasticity and Mode of Cell Migration. Dev Cell 45, 681–695.e684 (2018). 10.1016/j.devcel.2018.05.027

40 Lamouille, S., Xu, J. & Derynck, R. Molecular mechanisms of epithelial-mesenchymal transition. Nat Rev Mol Cell Biol 15, 178–196 (2014). 10.1038/nrm3758

41 Belvedere, R. et al. Role of intracellular and extracellular annexin A1 in migration and invasion of human pancreatic carcinoma cells. BMC Cancer 14, 961 (2014). 10.1186/1471-2407-14-961

42 Che, P. et al. S100A4 promotes pancreatic cancer progression through a dual signaling pathway mediated by Src and focal adhesion kinase. Sci Rep 5, 8453 (2015). 10.1038/srep08453

43 Pratap, A. et al. Glypican 1 promotes proliferation and migration in esophagogastric adenocarcinoma via activating AKT/GSK/β-catenin pathway. J Gastrointest Oncol 13, 2082–2104 (2022). 10.21037/jgo-22-240

44 Cordenonsi, M. et al. The Hippo transducer TAZ confers cancer stem cell-related traits on breast cancer cells. Cell 147, 759–772 (2011). 10.1016/j.cell.2011.09.048

45 Chan, S. W. et al. A role for TAZ in migration, invasion, and tumorigenesis of breast cancer cells. Cancer Res 68, 2592–2598 (2008). 10.1158/0008-5472.Can-07-2696

46 Islam, K. et al. Upregulated LAMA3 modulates proliferation, adhesion, migration and epithelial-to-mesenchymal transition of cholangiocarcinoma cells. Sci Rep 13, 22598 (2023). 10.1038/s41598-023-48798-8

47 Kirtonia, A. et al. Overexpression of laminin-5 gamma-2 promotes tumorigenesis of pancreatic ductal adenocarcinoma through EGFR/ERK1/2/AKT/mTOR cascade. Cell Mol Life Sci 79, 362 (2022). 10.1007/s00018-022-04392-1

48 Giannelli, G., Falk-Marzillier, J., Schiraldi, O., Stetler-Stevenson, W. G. & Quaranta, V. Induction of cell migration by matrix metalloprotease-2 cleavage of laminin-5. Science 277, 225–228 (1997). 10.1126/science.277.5323.225

49 Okada, Y., Takahashi, N., Takayama, T. & Goel, A. LAMC2 promotes cancer progression and gemcitabine resistance through modulation of EMT and ATP-binding cassette transporters in pancreatic ductal adenocarcinoma. Carcinogenesis 42, 546–556 (2021). 10.1093/carcin/bgab011

50 Wang, H., Cai, J., Du, S., Wei, W. & Shen, X. LAMC2 modulates the acidity of microenvironments to promote invasion and migration of pancreatic cancer cells via regulating AKT-dependent NHE1 activity. Exp Cell Res 391, 111984 (2020). 10.1016/j.yexcr.2020.111984

51 Moon, Y. W. et al. LAMC2 enhances the metastatic potential of lung adenocarcinoma. Cell Death Differ 22, 1341–1352 (2015). 10.1038/cdd.2014.228

52 Scialpi, R., Arrè, V., Giannelli, G. & Dituri, F. Laminin-332 γ2 Monomeric Chain Promotes Adhesion and Migration of Hepatocellular Carcinoma Cells. Cancers (Basel) 15 (2023). 10.3390/cancers15020373

53 Cañellas-Socias, A. et al. Metastatic recurrence in colorectal cancer arises from residual EMP1(+) cells. Nature 611, 603–613 (2022). 10.1038/s41586-022-05402-9

54 Escobar-Hoyos, L. F. et al. Keratin-17 Promotes p27KIP1 Nuclear Export and Degradation and Offers Potential Prognostic Utility. Cancer Res 75, 3650–3662 (2015). 10.1158/0008-5472.Can-15-0293

55 Liu, Z. et al. Keratin 17 activates AKT signalling and induces epithelial-mesenchymal transition in oesophageal squamous cell carcinoma. J Proteomics 211, 103557 (2020). 10.1016/j.jprot.2019.103557

56 Zhang, P. et al. Cytokeratin 17 activates AKT signaling to induce epithelial-mesenchymal transition and promote bladder cancer progression. BMC Urol 25, 77 (2025). 10.1186/s12894-025-01760-4

57 Wu, L., Ding, W., Wang, X., Li, X. & Yang, J. Interference KRT17 reverses doxorubicin resistance in triple-negative breast cancer cells by Wnt/β-catenin signaling pathway. Genes Genomics 45, 1329–1338 (2023). 10.1007/s13258-023-01437-y

