## Supplementary Files for "Keratin 17 Defines an Epithelial-Mesenchymal Transition-Associated Migratory Cell State in Pancreatic Ductal Adenocarcinoma"

**Supplementary Figure S1. Independent gene-set and single-cell correlation analyses confirm the *KRT17*-associated EMT/migration program.** (A) GSEA enrichment plot, Gene Ontology Biological Process term “Epithelial to Mesenchymal Transition.” (B) GSEA enrichment plot, Gene Ontology Biological Process term “Positive Regulation of Epithelial Cell Migration.” (C) Hallmark pathway enrichment (gene ratio) for *KRT17*-positive vs. *KRT17*-negative malignant cells. Asterisks denote pathways with significant enrichment (adjusted $p < 0.05$). (D) Expression prevalence of each marker, *KRT17*-positive vs. *KRT17*-negative cells. (E-J) Single-cell correlation of the markers with *KRT17* expression: ANXA1 (E, $\rho = 0.48$), S100A4 (F, $\rho = 0.37$), GPC1 (G, $\rho = 0.38$), LAMC2 (H, $\rho = 0.28$), WWTR1 (I, $\rho = 0.23$), LAMA3 (J, $\rho = 0.16$). (K-P) PC1/PC2 feature plots for ANXA1, S100A4, GPC1, LAMC2, WWTR1, and LAMA3.

**Supplementary Figure S2. Among the remaining five candidate markers, only GPC1 and LAMA3 show modest protein-level correlations with K17, and K17 and LAMC2 are each associated with shorter overall survival in the CPTAC cohort.** (A-E) Spearman correlation of K17 abundance with ANXA1 (A, $\rho = 0.111$, $p = 0.191$), S100A4 (B, $\rho = 0.135$, $p = 0.113$), GPC1 (C, $\rho = 0.291$, $p = 5.01 \times 10^{-4}$), WWTR1 (D, $\rho = -0.042$, $p = 0.619$), and LAMA3 (E, $\rho = 0.358$, $p = 1.61 \times 10^{-5}$) ($n = 140$ patients). Only GPC1 and LAMA3 reach significance. (F) Forest plot of single-marker (univariable) Cox models for overall survival: LAMC2-positive HR = 2.01 ($p = 0.004$) and K17-positive HR = 1.93 ($p = 0.009$). (G) Kaplan-Meier overall survival by K17 status ($n = 123$; K17-positive, $n = 53$; K17-negative, $n = 70$; HR = 1.93, $p = 0.009$). (H) Kaplan-Meier overall survival by LAMC2 status ($n = 123$; LAMC2-positive, $n = 72$; LAMC2-negative, $n = 51$; HR = 2.01, $p = 0.004$).

**Supplementary Figure S3. MACSima image analyses segment tumor cells and representative images confirm specific staining for every panel marker.** (A) Experimental workflow schematic. (B) Representative image showing the tumor-cell segmentation markers PanCK (white) and K17 (red). (C) Nuclear segmentation of tumor (yellow) and non-tumor (blue) cells. (D) Merged multiplex image. (E-L) Single-channel images confirming specific staining for PanCK (E), K17 (F), LAMC2 (G), ANXA1 (H), S100A4 (I), GPC1 (J), WWTR1 (K), and LAMA3 (L). Scale bars, 50 μm.

**Supplementary Figure S4. Among the remaining candidate markers, S100A4, WWTR1, and LAMA3 show no significant difference between K17-positive and K17-negative tumor cells, and GPC1 is enriched in K17-negative tumor cells.** (A-F) UMAP feature plots of the MACSima tumor-cell clusters, colored by marker expression (red, marker-positive; blue, marker-negative) for ANXA1 (A), S100A4 (B), LAMC2 (C), LAMA3 (D), GPC1 (E), and WWTR1 (F). Numbers indicate Leiden clusters. (G-J) Percentage of marker-positive cells among K17-positive versus K17-negative tumor cells for S100A4 (G; N = 68 cores; ns), GPC1 (H; N = 69 cores; \*\*\*\*p < 0.0001; higher in K17-negative tumor cells), WWTR1 (I; N = 69 cores; ns), and LAMA3 (J; N = 69 cores; ns). ns, not significant. Boxes show the median and interquartile range, whiskers show the range, and “+” marks the mean. Wilcoxon matched-pairs signed-rank test, with K17-positive and K17-negative tumor cells paired within each core. (K-N) Representative multiplex images showing PanCK (white) alone (K) and PanCK overlaid with K17 (red) (L), LAMC2 (green) (M), and ANXA1 (green) (N). Scale bars, 50 μm.

**Supplementary Figure S5. Single-plex immunohistochemistry confirms direct spatial overlap between K17 and LAMC2.** (A) K17 single-plex IHC (brown, DAB

chromogen), low magnification. (B) Higher-magnification inset of the boxed region in (A). (C) LAMC2 single-plex IHC (brown, DAB chromogen) performed on an adjacent serial section from the same tumor block as (A), demonstrating spatial overlap of K17 and LAMC2 staining within the same tumor architecture. (D) Higher-magnification inset of the boxed region in (C). Scale bars, 620 $\mu\text{m}$ (A, C) and 124 $\mu\text{m}$ (B, D).

**Supplementary Figure S6. K17 is expressed throughout the spheroid, and differences in proliferation do not readily account for its effect on migration.** (A-C) Confocal immunofluorescence for K17 (red) and Hoechst 33342 (blue) in whole-mount spheroids: L3.6 WT (A), L3.6 K17 (B), and PANC-1 K17 (C), showing K17 signal predominantly at the outer periphery. (D, E) K17 immunohistochemistry (brown) of FFPE sections from L3.6 EV (D) and L3.6 K17 (E) spheroids, showing K17 staining throughout the full depth of the spheroid, including the core, indicating that the peripherally restricted confocal signal in (A-C) reflects limited antibody/light penetration rather than true spatial restriction of K17 expression. (F-H) Ki-67 (green) immunofluorescence: L3.6 WT vs. KO (F), L3.6 EV vs. K17 (G), PANC-1 EV vs. K17 (H). (I) Quantification of Ki-67 index, L3.6 WT vs. KO (\*\*$p < 0.01$). (J) Quantification of Ki-67 index, L3.6 EV vs. K17-overexpressing (ns). (K) Quantification of Ki-67 index, PANC-1 EV vs. K17 (ns). ns, not significant. Scale bars, 400 $\mu\text{m}$ and 250 $\mu\text{m}$ for IF images, and 155 $\mu\text{m}$ for IHC images.

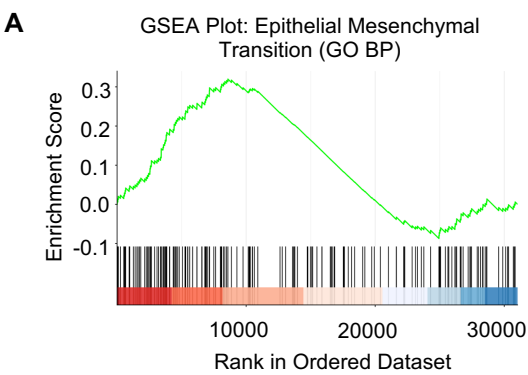

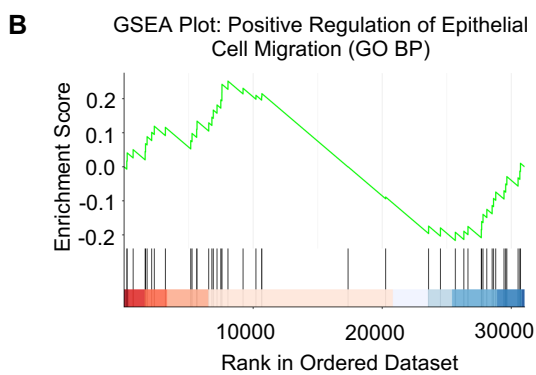

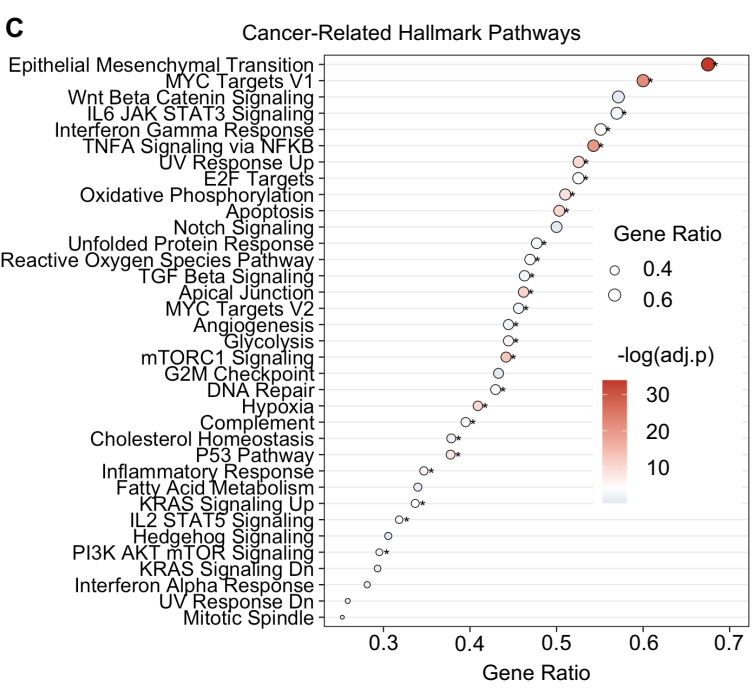

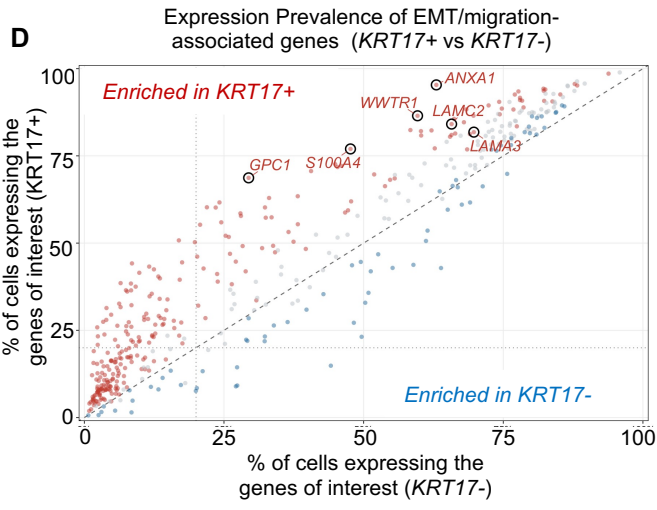

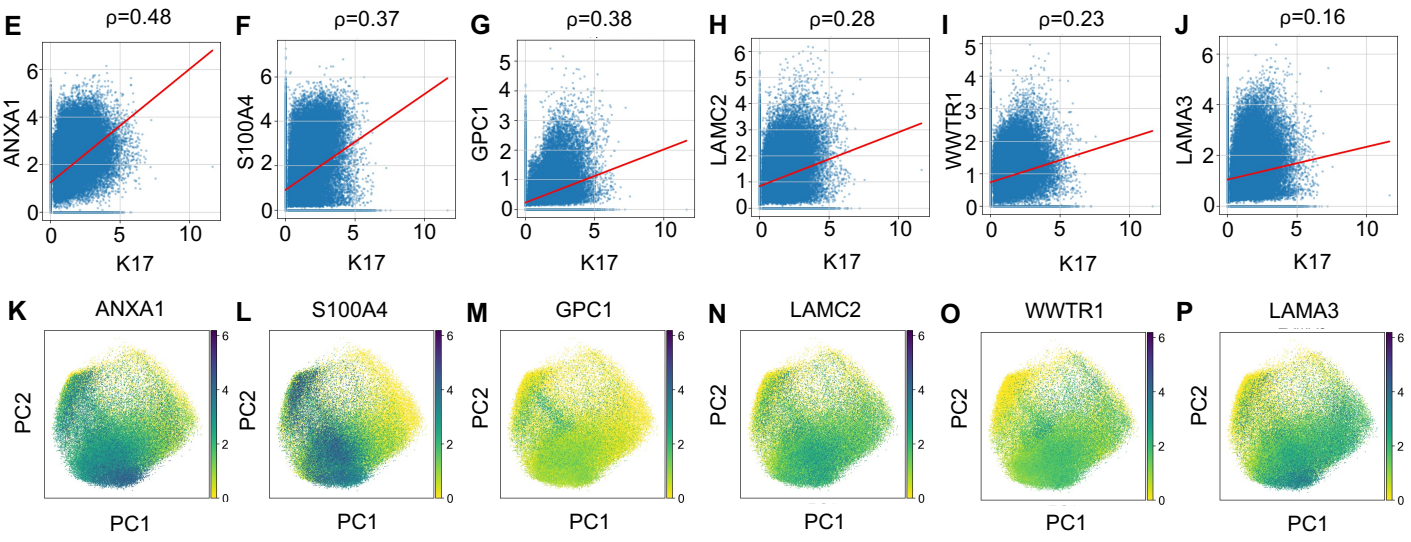

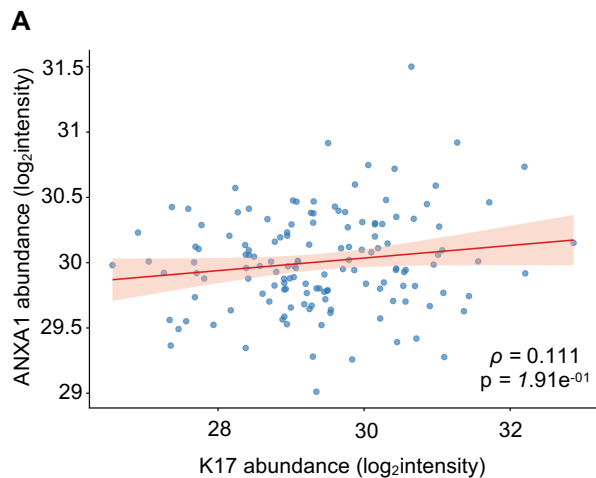

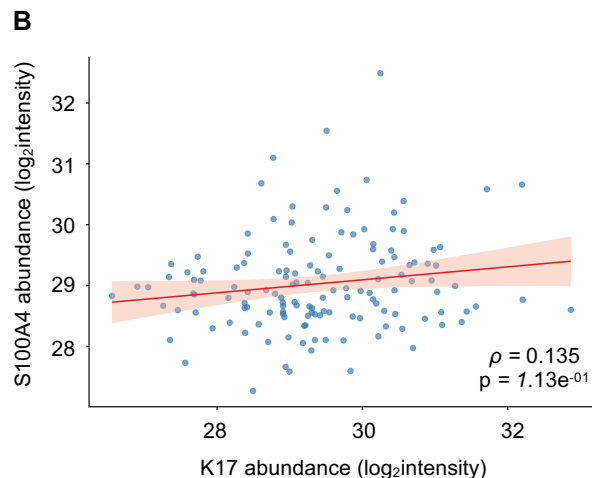

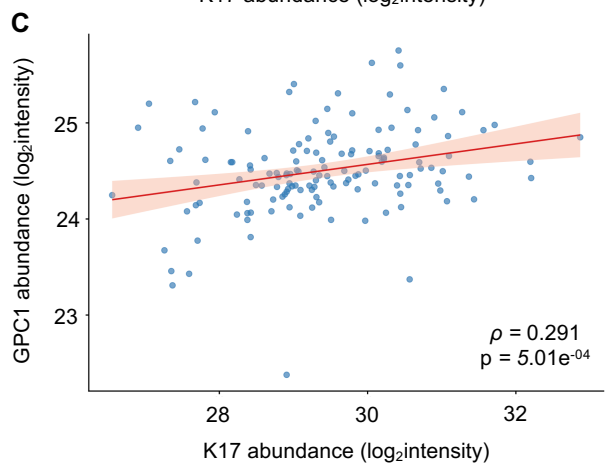

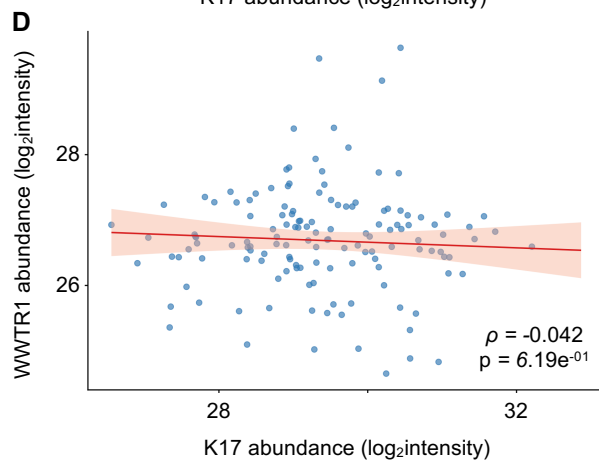

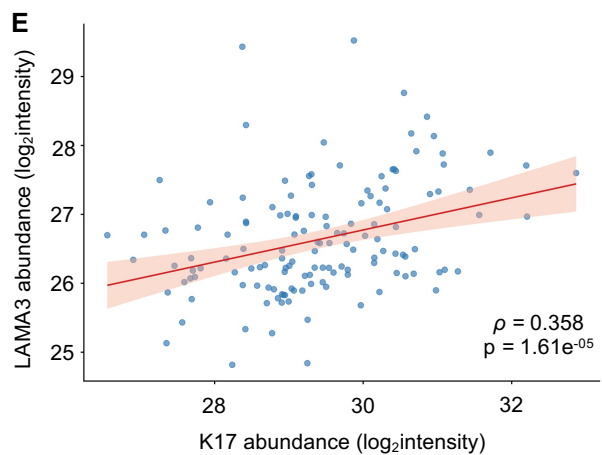

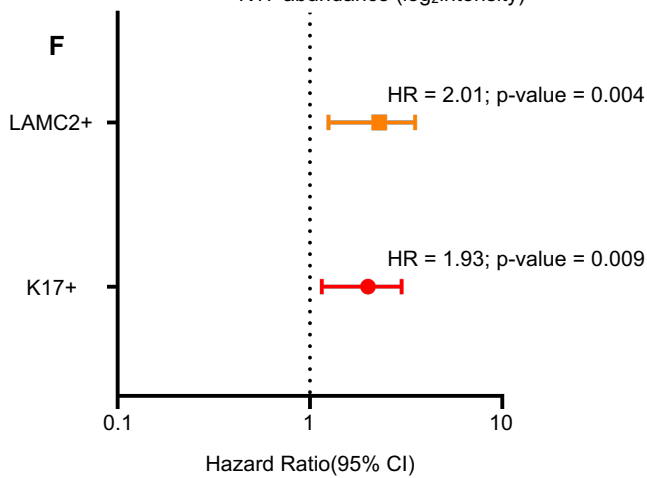

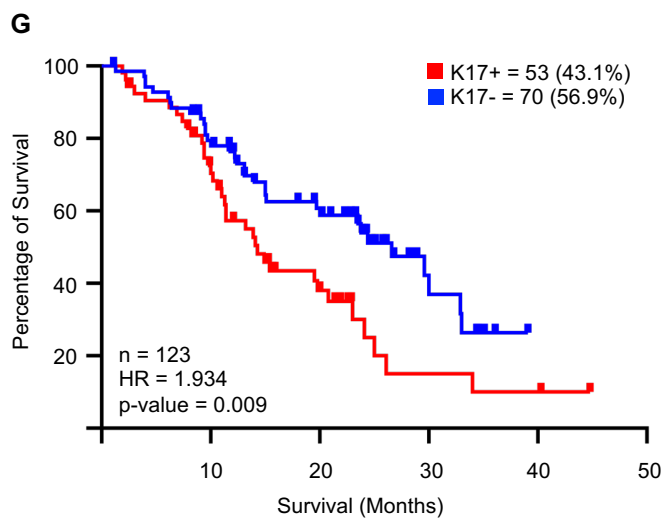

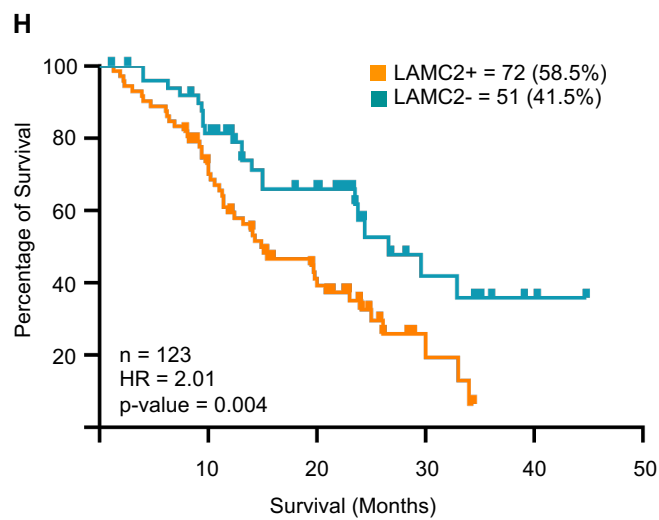

A

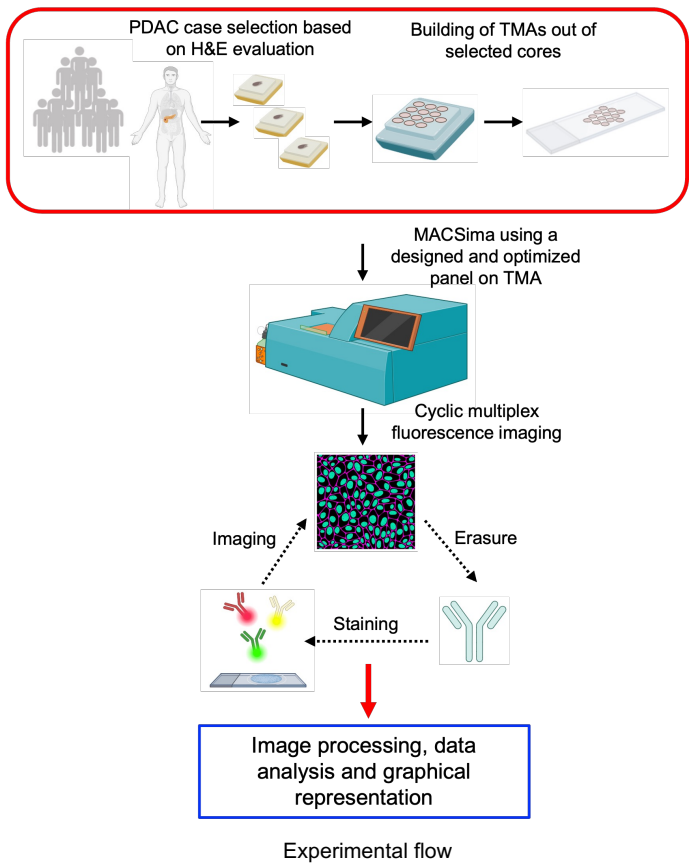

B

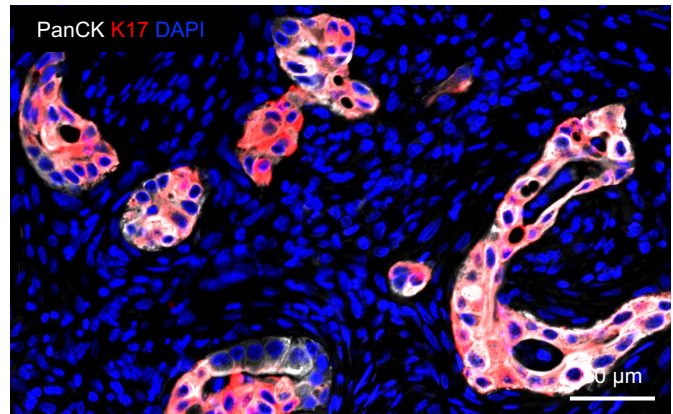

Representative image showing the segmentation markers (PanCK and K17)

C

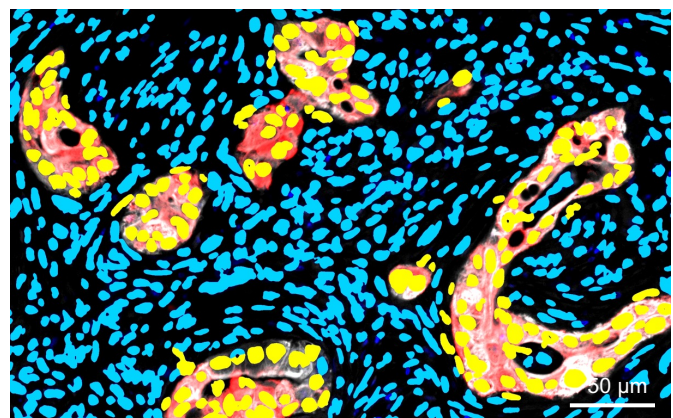

Nuclei detected of tumor (yellow) and non-tumor (blue) cells

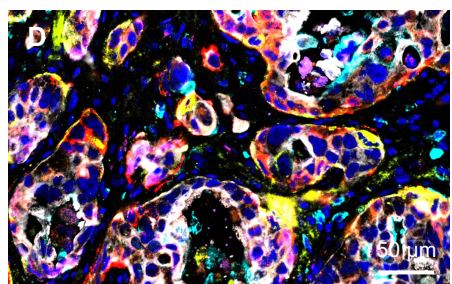

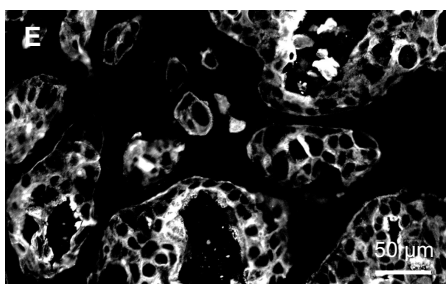

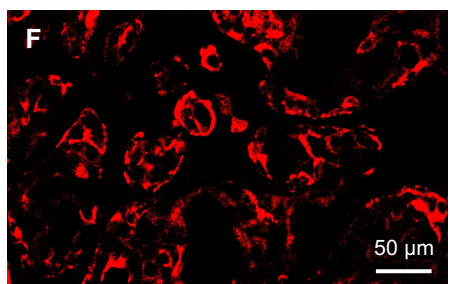

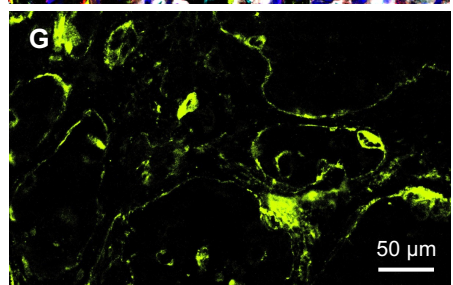

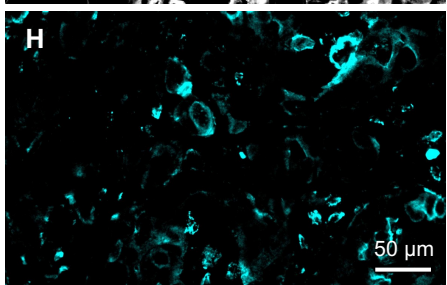

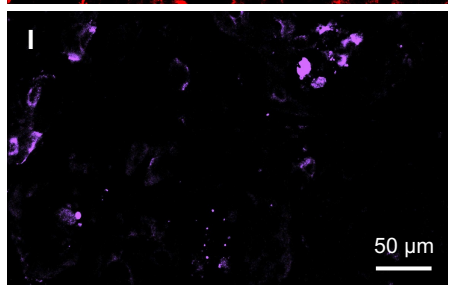

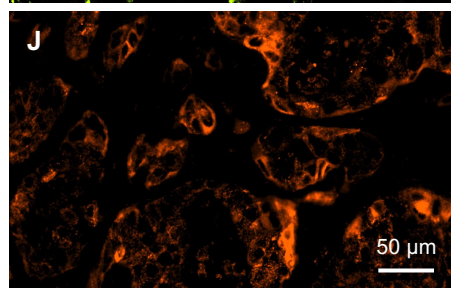

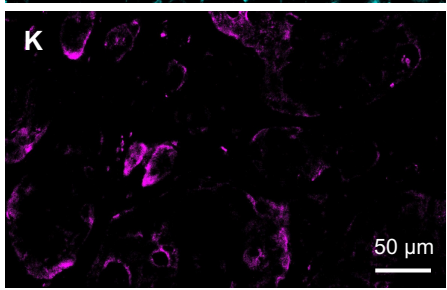

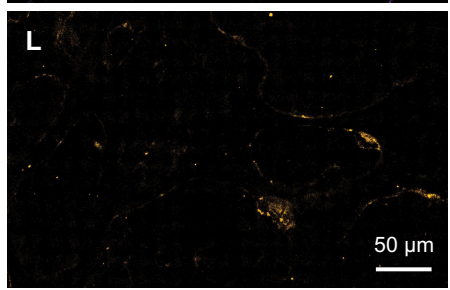

PanCK K17 LAMC2 ANXA1 S100A4 GPC1 WWTR1 LAMA3 DAPI

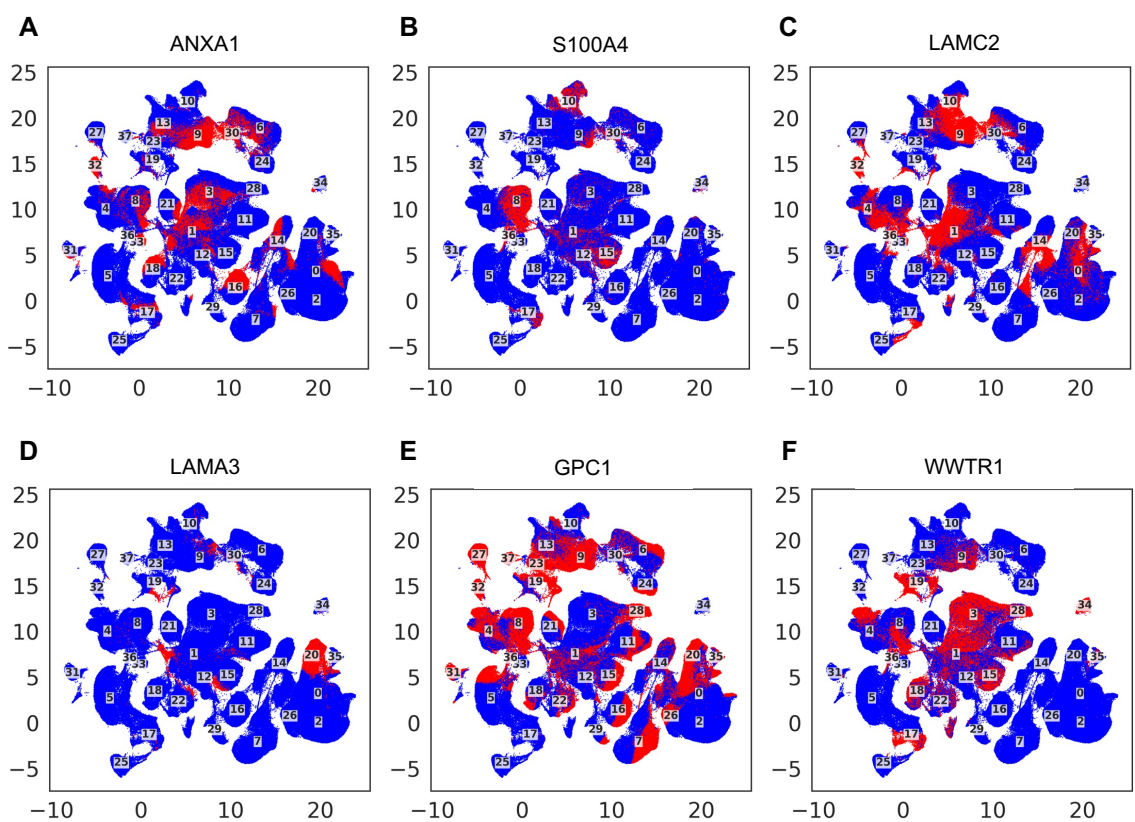

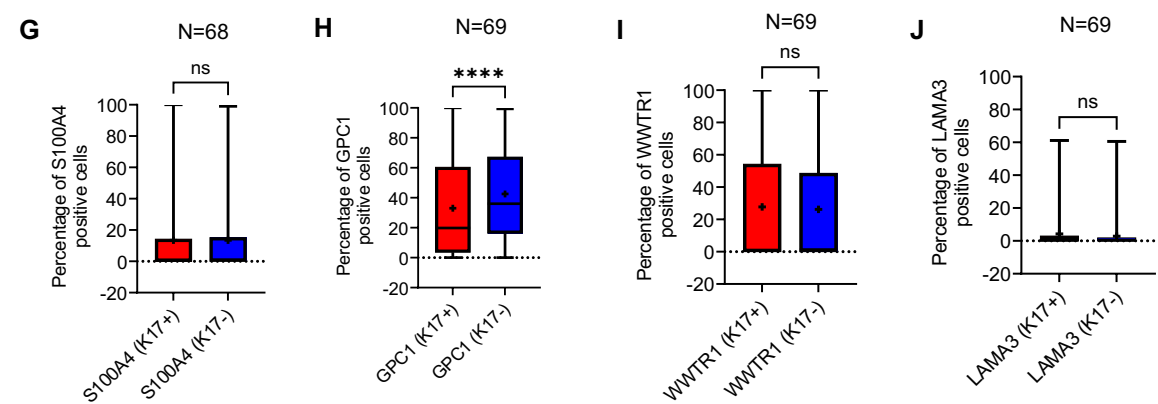

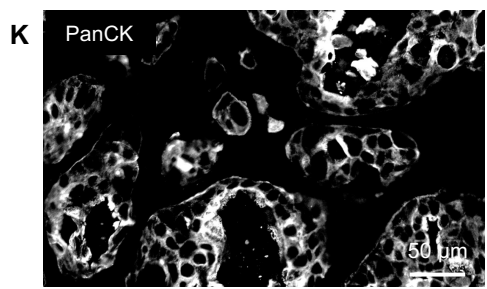

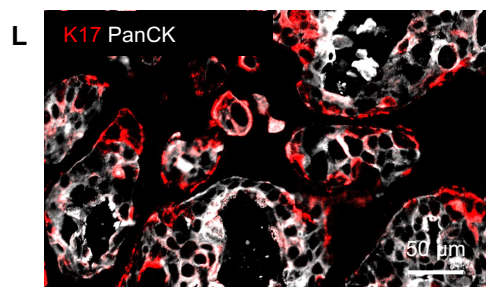

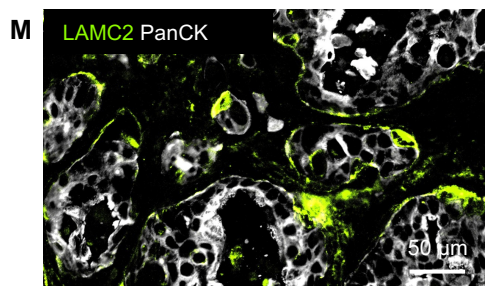

**Supplementary Table S1. Demographic and Clinicopathologic Characteristics of the MACSima Cohort**

| **Characteristic** | **n** | **%** |
| --- | --- | --- |
| Total unique patients | 63 |  |
| Total tissue cores profiled | 69 |  |
| **Age at diagnosis, years** |  |  |
| Mean ± SD | 69.8 ± 10.2 |  |
| Median (range) | 72 (43–89) |  |
| **Sex** |  |  |
| Female | 34 | 54.0% |
| Male | 29 | 46.0% |
| **Race** |  |  |
| White | 39 | 61.9% |
| Black | 3 | 4.8% |
| Other | 3 | 4.8% |
| Not available | 18 | 28.6% |
| **Vital status** |  |  |
| Dead | 32 | 50.8% |
| Alive | 13 | 20.6% |
| Not available | 18 | 28.6% |
| **Pathologic AJCC stage group (pre-2018)** |  |  |
| Stage 1A | 2 | 3.2% |
| Stage 1B | 7 | 11.1% |
| Stage 2A | 11 | 17.5% |
| Stage 2B | 30 | 47.6% |
| Stage 3 | 11 | 17.5% |
| Stage 4 | 2 | 3.2% |

**Supplementary Table S2. MACSima Multiplex Antibody Panel**

| Marker | Vendor | Catalog Number | Clone | Fluorophore | RRID | Dilution | Panel Status |
| --- | --- | --- | --- | --- | --- | --- | --- |
| Cytokeratin LMW (PanCK) | Miltenyi Biotec | 130-112-743 | REA831 | FITC | RRID:AB_2651496 | 1:50 | Included |
| Cytokeratin 17 (K17) | Cell Signaling Technology | 12509 | D12E5 | PE | RRID:AB_2797939 | 1:50 | Included |
| Annexin A1 (ANXA1) | Miltenyi Biotec | 130-119-354 | REA1122 | APC | RRID:AB_2857450 | 1:50 | Included |
| S100A4 | Miltenyi Biotec | 130-134-196 | REA1572 | Vio B515 | RRID:AB_3664964 | 1:50 | Included |
| GPC1 | Abcam | ab237290 | EPR19285 | Alexa Fluor 647 | Not available | 1:50 | Included |
| LAMC2 | Santa Cruz Biotechnology | sc-28330 AF647 | E-6 | Alexa Fluor 647 | RRID:AB_2134476 | 1:50 | Included |
| WWTR1 | Proteintech | CL488-23306 |  | CoraLite Plus 488 | RRID:AB_3672754 | 1:50 | Included |
| LAMA3 | Novus Biologicals | NBP3-44416AF647 | CL3112 | Alexa Fluor 647 | Not available | 1:50 | Included |

**Supplementary Table S3. MACSima Segmentation Parameters**

| **Parameter** | **Value / Setting** |
| --- | --- |
| **Nucleus** |  |
| Method | Advanced Morphology for Tissue |
| Reference Channel | DAPI C1 |
| Min/Max Diameter | 25 / 120 pixels |
| Detection Sensitivity | 200% |
| Separation Force | 90% |
| Smoothing Filter Sigma | 3.0 |
| **Cytoplasm** |  |
| Active | Yes |
| Method | Constrained Donut |
| Constraint Channels | Multiple Channels Selected – Cytokeratin LMW and Keratin 17 |
| Detection Sensitivity | 1000% |
| Donut Width | 100 pixels |
| **Measure Cell Features** |  |
| Active | Yes |
| All Biomarker Expressions | Yes |
| Cell and Nuclear Shape | Yes |
| Measure Channels | Automatic |
| Export Nuclei Labels | None |
| Export Cell Labels | None |
| **General Settings** |  |
| Calibrated Distance Units | No |
| Log Diagnostics | Yes |
| Remove Edge Objects | No |
| Use Exclusion Regions | Yes |
